# A genome-wide CRISPRi screen reveals a StkP-mediated connection between cell-wall integrity and competence in *Streptococcus salivarius*

**DOI:** 10.1101/2022.06.29.498087

**Authors:** Adrien Knoops, Alexandra Waegemans, Morgane Lamontagne, Baptiste Decat, Johann Mignolet, Jan-Willem Veening, Pascal Hols

## Abstract

Competence is one of the most efficient bacterial evolutionary and adaptative strategies by synchronizing production of antibacterial compounds and integration of DNA released by dead cells. In most streptococci, this tactic is orchestrated by the ComRS system, a pheromone communication device providing a sharp time window of activation in which only part of the population is responsive. Understanding how this developmental process integrates multiple inputs to fine-tune the adequate response is a long-standing question. However, essential genes involved in the regulation of ComRS have been challenging to study. In this work, we built a conditional mutant library using CRISPR-interference and performed three complementary screens to investigate competence genetic regulation in the human commensal *Streptococus salivarius*. We show that initiation of competence increases upon cell-wall impairment, suggesting a connection between cell envelope stress and competence activation. Notably, we report a key role for StkP, a serine-threonine kinase known to regulate cell-wall homeostasis. We show that StkP controls competence by a mechanism that reacts to peptidoglycan fragments. Together, our data suggest a key cell-wall sensing mechanism coupling competence to cell envelope integrity.

**IMPORTANCE:** Survival of human commensal streptococci in the digestive tract requires efficient strategies which must be tightly and collectively controlled for responding to competitive pressure and drastic environmental changes. In this context, the autocrine signaling system ComRS controlling competence for natural transformation and predation in salivarius streptococci could be seen as a multi-input device integrating a variety of environmental stimuli. In this work, we revealed novel positive and negative competence modulators by using a genome-wide CRISPR-interference strategy. Notably, we highlighted an unexpected connection between bacterial envelope integrity and competence activation that involves several cell-wall sensors. Together, these results showcase how commensal streptococci can fine-tune the pheromone-based competence system by responding to multiple inputs affecting their physiological status in order to calibrate an appropriate collective behavior.

## INTRODUCTION

In the human digestive tract, bacteria face highly competitive pressure and physicochemical challenges. Surviving in this environment requires powerful and efficient strategies which must be tightly controlled and collectively coordinated (1–3). Quorum sensing (QS) devices are particularly suited to control concerted survival tactics since they perform bacterial density sensing. Initially thought to be restricted to this role, recent evidences suggest that QS systems can operate as autocrine modules and process multiple inputs (4). On the one hand, QS autocrine signaling allows heterogeneity amplification by positive feedback loops, a key feature for sub-populations activation (5–7). On the other hand, environmental stimuli can fine-tune the sensitivity of the pheromone-based apparatus (8, 9). This latter property is switching the QS property from a cell-density to a multi-input device, integrating diverse stimuli to calibrate population-wide strategies (4).

One of the best characterized QS-mediated process in Gram-positive bacteria is competence regulation (10). Orchestrating predation through bacteriocin production together with natural transformation, competence is regulated by two types of signaling systems in streptococci (11). The ComCDE system found in the mitis and anginosus groups relies on the sensing of the extracellular pheromone CSP (<u>c</u>ompetence <u>s</u>timulating <u>p</u>eptide) that induces a phosphorelay leading to transcriptional activation of competence genes comprising *comX*, which codes for the master competence-specific sigma factor (12). The alternative predominant system in streptococci is based on the production/maturation of the pheromone XIP (*com<u>X</u>*-inducer <u>p</u>eptide), which is internalized by the Opp transporter and binds the intracellular receptor ComR (13, 14). Subsequently, the dimeric ComR**·**XIP complex activates several bacteriocin and competence genes including *comX* (15–17).

Uncovering the environmental triggers allowing permissive conditions for competence QS has remained challenging in streptococci (18). Since two-component systems (TCS) and serine-threonine kinases (STK) are dedicated to sense the outside world, they constitute attractive targets to couple environmental stimuli to QS reactivity. In *Streptococcus pneumoniae,* several of those sensors (e.g. StkP, CiaRH, VicRK) have been highlighted to control the ComCDE activity upon pH, O_2_, cell density or antibiotic stresses (9, 19–23). In the cariogenic *Streptococcus mutans* species, other distal regulators have been highlighted such as ScnRK, HdrM, BrsRM, CiaRH or StkP that link competence activation to various growth conditions (pH, carbohydrate source, oxygen, cell density) (24–31). In salivarius streptococci, we recently uncovered a regulatory inhibition of the CovRS environmental sensor on the ComRS signaling system (7). As exemplified by these three cases, despite the fact that environmental triggers can be shared, environmental sensors bridging detection of stimuli to competence can be highly divergent between species.

To investigate key sensors generating permissive conditions for competence activation, genome-wide screens are the fastest and most-suited approaches. While Tn-seq strategies have already revealed several regulators in *S. mutans* and *S. pneumoniae* (32, 33), classical knock- out characterization of the identified genes is often impaired by their essentiality. Recently, a genome-wide CRISPR-interference (CRISPRi) screening method was shown to overcome this drawback for *Escherichia coli* and *S. pneumoniae* (34–36). This technique combines the use of a guide RNA (gRNA) library targeting the whole genome together with a catalytically dead mutant of Cas9 (dCas9), producing transcriptional interference upon DNA binding. Plugging the dCas9 under the control of an inducible promoter allows the construction of a conditional mutant library which can be used for genetic screens and further for characterization of essential genes by knocking down their expression (34, 35).

In this work, we used this technique in combination with three distinct screens to unveil novel competence regulators. Cross-validation of the hits obtained from the three screens converged towards a connection between impairment of cell-wall biogenesis and competence activation.

Coherently, several sensors of the bacterial envelope integrity were identified among which StkP, suggesting a putative signaling pathway bridging cell-wall stress to competence activation.

## RESULTS

### Screening for spontaneous transformation by genome-wide CRISPRi inhibition

To identify unknown modulators of competence in *S. salivarius* HSISS4 (37), we set up a genome-wide CRISPRi strategy. To design gRNAs on the whole genome of HSISS4, we subset all the 20 nt sequences followed by a protospacer adjacent motif (PAM, NGG sequence) and retained only gRNAs binding the coding strand (non-template strand) of coding sequences (CDSs) and both strands in intergenic regions (34). We ended up with a total of 83,103 gRNAs (Table S1A) that were introduced under the control of a constitutive promoter (P_3_, (38)) at a neutral chromosomal locus. The random chromosomal distribution of gRNAs in the library was preliminary evaluated by the direct sequencing of 40 individual clones (Fig. S1A).

The transfer of the library was initially performed in a strain carrying an IPTG-inducible dCas9 (P_F6_*-lacI;* P*_lac_-dcas9*, (35)), which was previously validated for generating CRISPRi conditional mutants (7) (Fig. 1). To evaluate the functionality of the library, this first strain was screened for the activation of spontaneous natural transformation. We hypothesized that dCas9-mediated repression of genes involved in competence inhibition (i.e. antagonist genes) will result in spontaneous natural transformation and donor DNA integration. We activated the interference library by adding IPTG (dCas9 activation) to a liquid culture supplemented with donor DNA containing a chloramphenicol resistance cassette (Fig. 1A). We were able to isolate 16 candidates after 3 independent rounds of selection, all harboring a different gRNA (Table 1). In order to confirm the phenotype generated by these gRNAs, we back-transformed them individually into the original strain and assessed their transformability. Spontaneous transformation was confirmed for 10 candidates (Table 1). Importantly, this functional screen succeeded to identify two previously described negative effectors of competence acting on ComX or XIP stability (*clpC* and *pepF*, respectively) (39, 40).

**Figure 1.**
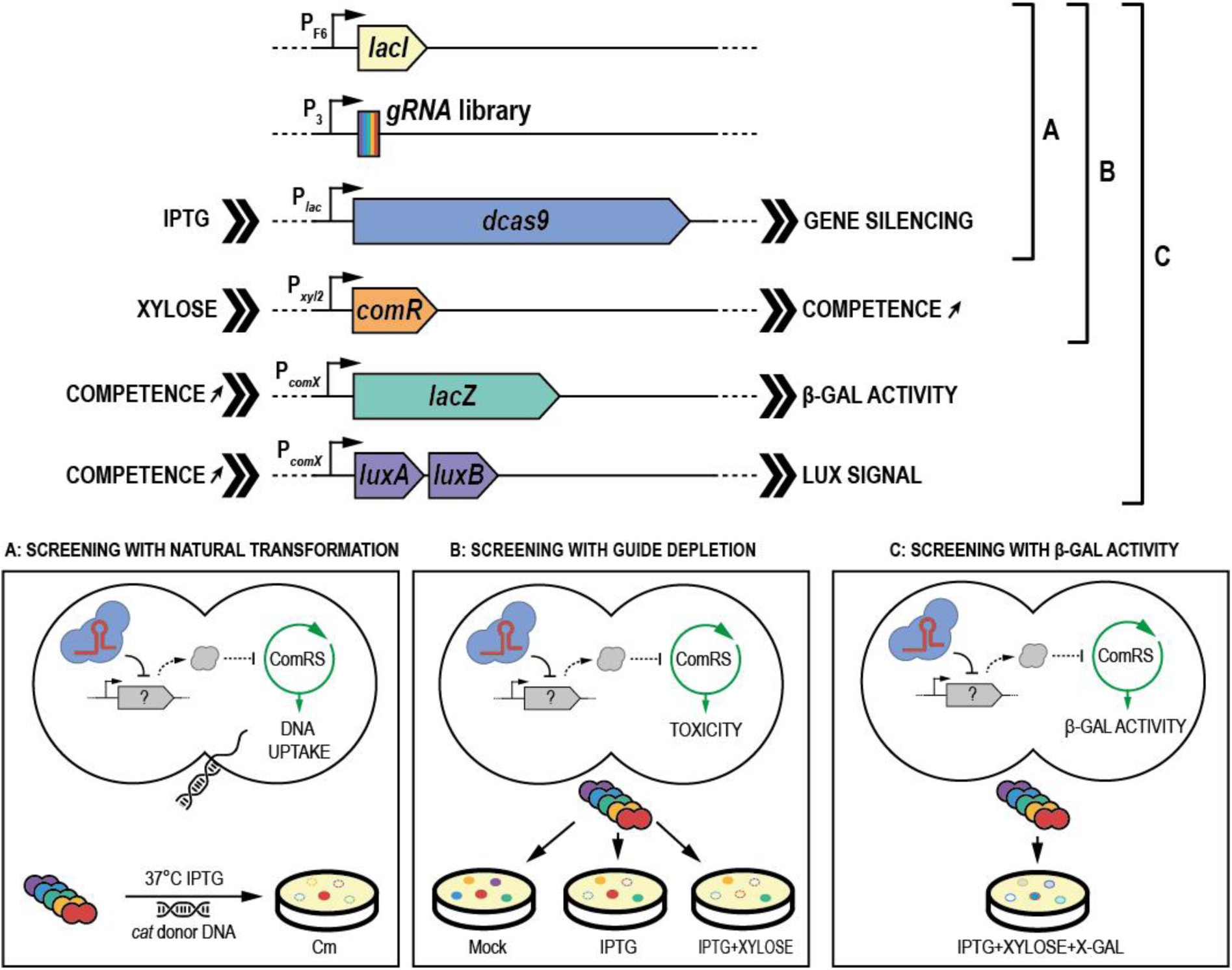
CRISPRi screening strategies for competence modulators in *S. salivarius.* A library of gRNAs was designed and introduced (P_3_*-gRNA*) in an engineered strain of *S. salivarius* harboring an IPTG-inducible system for dCas9 (P_F6_*-lacI*; P*_lac_-dcas9*). A first library was screened for spontaneous competence activation upon dCas9-inhibition by growing cells in chemically defined medium in presence of IPTG and *cat* donnor DNA. The selection on chloramphenicol plates was associated with inhibition of competence negative players (**A**). A second library was generated by introducing the gRNA library into the same background with a supplemental construct consisting of a xylose-inducible promoter fused to *comR* (P*_xyl2_-comR*). The library was spread on control (mock), gRNA library-induced (IPTG) or gRNA library- and competence-induced (IPTG + xylose) plates. NGS analysis of depleted gRNAs in the three conditions was performed to search for costly genes only associated to competence (**B**). A third library was built by adding *lacZ* under the control of P*_comX_* (P*_comX_-lacZ*) together with a competence luciferase reporter system (P*_comX_*-*luxAB*) to the previous strain and transferring the gRNA library into this background. The generated library was screened on plates containing IPTG, xylose and X-gal. gRNAs targeting potential competence inhibitory or activatory genes were associated to dark blue or white phenotypes, respectively (**C**).

**Table 1.**
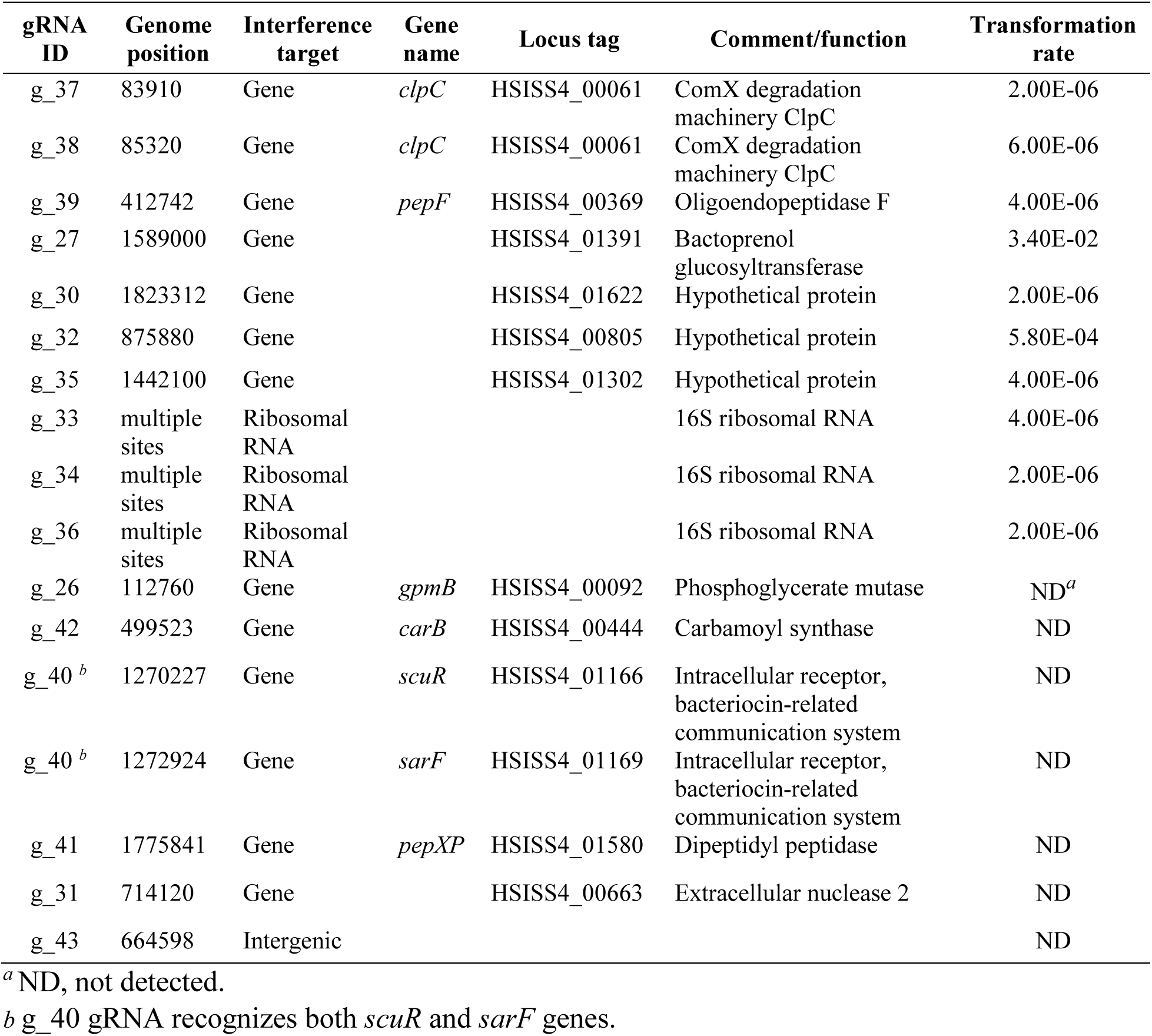
gRNA identification in spontaneous transformants.

### Screening based on gRNA depletion

This strategy was based on the toxicity of competence overactivation in the strain HSISS4 (16). We assumed that repression of competence-antagonist genes would produce a fitness cost, resulting in pool depletion of gRNAs targeting the corresponding genes. To set up this strategy, a second screening strain was generated by introducing a supplemental construct consisting of a xylose-inducible *comR* gene (P*_xyl2_*-*comR*), allowing a mild competence activation upon addition of xylose, a non-metabolizable sugar in *S. salivarius* (7) (Fig. 1B). After introducing the gRNA library into this strain, we spread it on three different solid culture conditions. The first condition without any inducer was used as control (mock). The second condition was induced with IPTG alone to activate the CRISPRi library (Ci) and the third condition was supplemented with IPTG and xylose to concomitantly activate the CRISPRi library and competence (Ci+C). We hypothesized that we could identify modulator genes of competence by comparing the depletion of gRNAs between conditions Ci and Ci+C. To this aim, we performed high-throughput next-generation sequencing (NGS) to quantify each gRNA abundance per condition (Table S1). We first evaluated the randomness and homogeneity of gRNA distribution without any selection pressure (mock) by visualizing the mapping of the gRNAs on the genome of HSISS4 (Fig. S1B). Validating our previous Sanger-sequencing data (Fig. S1A), we showed that 99.7% (82,864 out of 83,104) of gRNAs were cloned in the non-induced library with an unbiased distribution (Table S1B and Fig. S1C). We next used the MAGeCK algorithm (41) to compare depletion of gRNAs between two conditions. As expected, the analysis of gRNA depletion between Ci and mock conditions uncovered well-known essential genes in streptococci (Table S2, Fig. S2A), as well as competence-related genes (e.g. *covR*, *pepF*) whose inactivation was recently shown to be lethal in strain HSISS4 (7, 40). We next performed the same analysis for the comparison of condition Ci+C with the mock (Table S3, Fig. S2B) and plotted against each other the two scores obtained from the two comparisons (i.e. Ci vs mock and Ci+C vs mock) (Fig. S3). As expected, depletion scores in the two comparisons displayed a high correlation showing that gene fitness (i.e. positive, neutral or negative) was conserved with or without competence activation (linear regression of R²=0.97). However, several outliers were present. Because they represent genes differentially affected in-between two conditions, we analyzed the standardized residuals of the linear regression (Fig. 2A, Table S4). We set up an arbitrary cut-off at +2.5 and -2.5 to identify the most representative outliers. Several depleted gRNAs were found as targeting genes known as competence antagonists such as the *mecA* gene encoding the ComX adaptor of the Clp degradation machinery (standardized residuals < -2.5, Table 2) (42). Unexpectedly, many crucial genes for competence activation (*comR*, *amiACDEF*) or competence-based bacteriocin production/immunity (e.g. *slvX-HSISS4_01664* operon) also showed gRNA depletion (Table 2) (16). In the strain HSISS4, competence, bacteriocins, and bacteriocin-immunity genes are concomitantly activated through ComR (16). Therefore, those genes might have been selected because a reduced competence activation goes along with a lower immunity rate toward bacteriocins, ultimately leading to a high fitness cost. Indeed, since bacteriocin producers are present at high cell density on plates due to xylose-mediated competence activation, non-competent and immunity-deficient cells will be killed through the well-established fratricide process (43).

**Figure 2.**
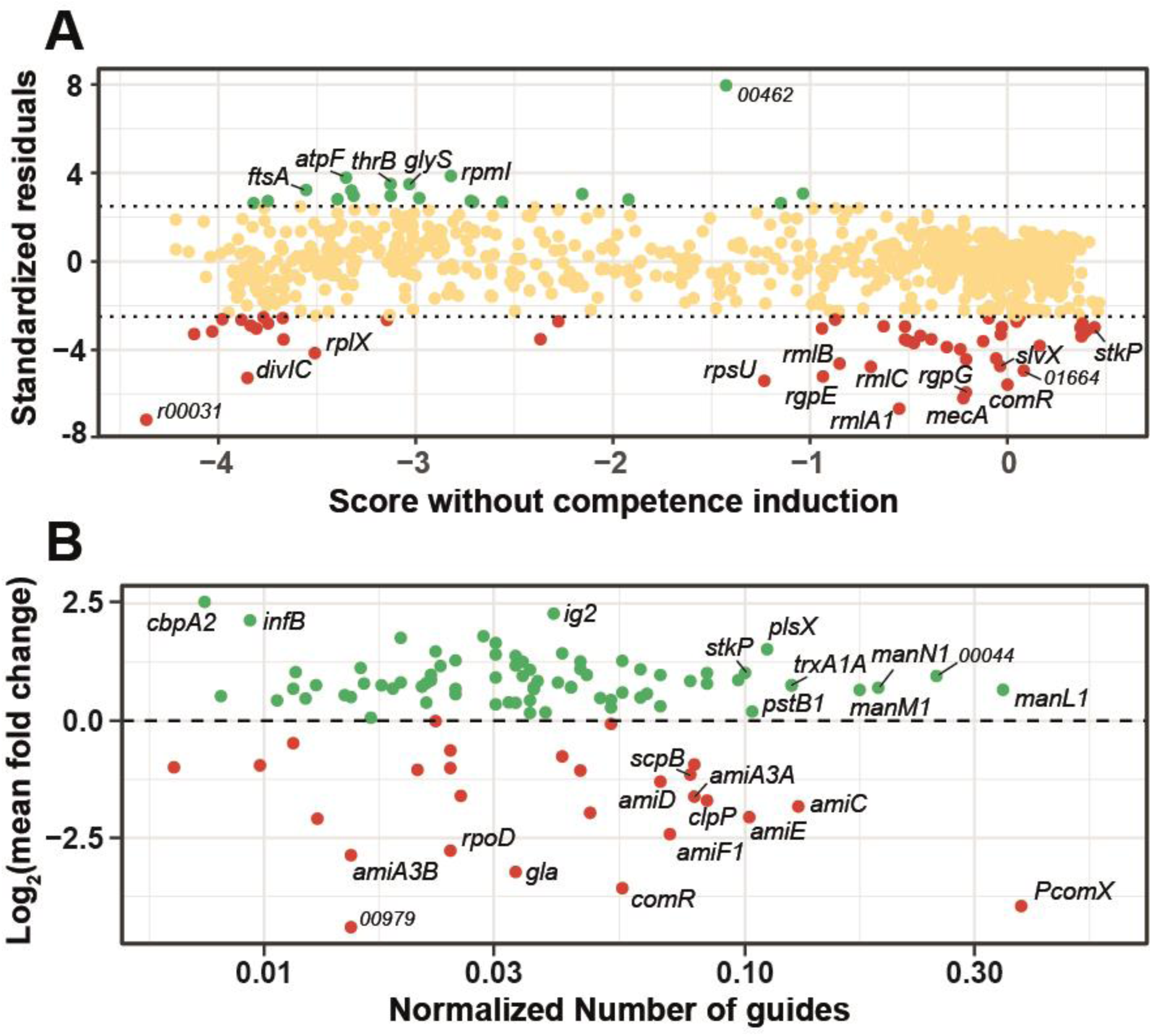
Selection of genes from CRISPRi screens. **A.** gRNA depletion screen. The gRNA library was grown on M17G plates for ∼ 12 generations with no induction (mock), with gRNA library induction (Ci), or with gRNA library and competence induction (Ci+C). The gRNAs (4 technical replicates per condition, ∼40 M reads) were sequenced and mapped thanks to the MAGeCK-VISPR algorithm (41). Using the same tool, we identify gRNA depletion in costly genes linked to library induction only (Ci vs mock) and both library and competence induction (Ci vs Ci+C) (Fig. S2). We then compared the gRNA depletion scores for each gene in both induction systems and performed a linear regression (Fig. S3). Standardized residuals of the regression were then computed and plotted in function of the score of each gene in the condition without competence induction (Ci). Positive (green) and negative (red) standardized residuals (arbitrary cut-off of +2.5 and -2.5) denote genes with enriched or depleted gRNAs, respectively. Dots in yellow are considered as non-significantly affected genes. **B.** β-Gal screen. After library production (∼10^5^ colonies), screening for dark blue and white clones on M17GL plates (with IPTG, xylose, and X-gal; P*_comX_-lacZ*, P*_xyl2_*-*comR*) and validation with luciferase assays (P*_comX_- luxAB*), clones with the most dissimilar luciferase phenotypes (141 dark blue and 68 white clones) were sequenced for gRNA identification. The y-axis displays the mean fold change Log_2_ value of luciferase activity calculated on all the gRNAs targeting the same gene. The x- axis displays the number of gRNAs targeting the same gene normalized by the expected total number of gRNAs present in the library for this gene. Green dots and red dots denote gene inhibition resulting in competence overactivation or repression, respectively.

**Table 2.** Identification of competence costly genes from the gRNA depletion screen.

| Gene name | Locus tag | Comment/function | Fitness cost score without competence induction <sup>a</sup> | Std residual (< -2.5) <sup>b</sup> |
| --- | --- | --- | --- | --- |
| **Competence-related** |  |  |  |  |
| *comR* | HSISS4_00217 | Competence intracellular receptor | 0.00 | -5.58 |
| *amiF1* | HSISS4_01361 | Oligopeptide ABC transporter, ATP-binding subunit F | -0.52 | -2.94 |
| *amiE* | HSISS4_01362 | Oligopeptide ABC transporter, ATP-binding subunit E | -0.47 | -3.68 |
| *amiD* | HSISS4_01363 | Oligopeptide ABC transporter, permease subunit D | -0.52 | -3.53 |
| *amiC* | HSISS4_01364 | Oligopeptide ABC transporter, permease subunit C | -0.51 | -3.59 |
| *amiA3A* | HSISS4_01365 | Oligopeptide ABC transporter, oligopeptide binding subunit A | -0.44 | -3.37 |
|  | HSISS4_01664 | SlvX immunity protein | 0.08 | -4.94 |
| *slvX* | HSISS4_01665 | Bacteriocin | -0.04 | -4.73 |
| *mecA* | HSISS4_00128 | ComX degradation machinery adaptor protein | -0.22 | -6.19 |
| *spxA1* | HSISS4_00943 | Transcriptional regulator | 0.38 | -2.80 |
| **Cell-envelope-related** |  |  |  |  |
| *stkP* | HSISS4_01348 | Serine-Threonine kinase | 0.44 | -2.98 |
| *acpP1* | HSISS4_00021 | Acyl carrier protein | 0.06 | -2.54 |
| *rgpG* | HSISS4_00129 | Polysaccharide synthesis protein | -0.21 | -5.92 |
| *rgpF* | HSISS4_01378 | Polysaccharide synthesis protein | -0.21 | -4.43 |
| *rgpE* | HSISS4_01379 | Extracellular rhamnan synthesis protein | -0.94 | -5.20 |
| *rgpA2* | HSISS4_01383 | Extracellular rhamnan synthesis protein | -2.37 | -3.52 |
| *rmlA1* | HSISS4_00723 | Rhamnose synthesis protein | -0.55 | -6.66 |
| *rmlC* | HSISS4_00724 | Rhamnose synthesis protein | -0.69 | -4.77 |
| *rmlB* | HSISS4_00725 | Rhamnose synthesis protein | -0.85 | -4.62 |
| *pgmA* | HSISS4_01102 | Phosphoglucomutase | -0.12 | -3.61 |
| *dgk* | HSISS4_00536 | Lipid carrier recycler | -2.28 | -2.71 |
| *murG* | HSISS4_00684 | Peptidoglycan Lipid II precursor synthesis | -3.98 | -2.62 |
|  | HSISS4_00889 | Exporter of O-antigen, teichoic acids, lipoteichoic acids (WpsG) | -0.24 | -3.97 |
| *dltD* | HSISS4_01108 | Poly(glycerolphosphate chain) D-alanine transfer protein | -0.03 | -2.98 |
| *dltC* | HSISS4_01109 | D-Alanine phosphoribitol ligase subunit 2 | -0.48 | -3.68 |
| *dltB* | HSISS4_01110 | D-alanyl transfer protein | -0.31 | -3.87 |
| *dltA* | HSISS4_01111 | D-Alanine phosphoribitol ligase subunit 1 | -0.39 | -3.52 |
| *dltX* | HSISS4_01112 | D-Ala-teichoic acid biosynthesis protein | -0.63 | -2.93 |
| *pstB1* | HSISS4_00936 | Phosphate transport, ATP binding protein | 0.37 | -3.00 |
| *pstC2* | HSISS4_00937 | Phosphate transport, permease protein | 0.38 | -3.39 |
| *pstC1* | HSISS4_00938 | Phosphate transport, permease protein | 0.38 | -3.27 |
| *pstS* | HSISS4_00939 | Phosphate transport, phosphate binding protein | 0.40 | -3.14 |
| *divIC* | HSISS4_00008 | Cell division protein | -3.85 | -5.27 |
| *ftsL* | HSISS4_01598 | Cell division protein | -4.12 | -3.29 |
| **Translation** |  |  |  |  |
| *prfB* | HSISS4_00848 | Peptide chain release factor | -0.06 | -4.39 |
| *proS* | HSISS4_00152 | Prolyl tRNA synthetase | -3.77 | -2.53 |
| *rplM* | HSISS4_00076 | Large subunit ribosomal protein | -3.67 | -3.53 |
| *rplX* | HSISS4_01812 | Large subunit ribosomal protein | -3.51 | -4.15 |
| *rpsU* | HSISS4_01396 | Small subunit ribosomal protein | -1.23 | -5.40 |
| *rpsF* | HSISS4_01661 | Small subunit ribosomal protein | -3.81 | -3.04 |
| *rpsE* | HSISS4_01806 | Small subunit ribosomal protein | -3.88 | -2.64 |
| *rpsS* | HSISS4_01819 | Small subunit ribosomal protein | -3.84 | -2.90 |
|  | HSISS4_00271 | Ribosomal protein | -3.75 | -2.80 |
|  | HSISS4_r00031 | tRNA Met | -4.37 | -7.16 |
|  | HSISS4_r00059 | tRNA Glu | -4.03 | -3.17 |
|  | HSISS4_r00070 | tRNA Arg | -3.67 | -2.56 |
| **Others** |  |  |  |  |
| *ctsR* | HSISS4_00060 | Stress transcriptional regulator | -0.09 | -2.57 |
| *atpE* | HSISS4_00399 | ATP synthase | -3.15 | -2.65 |
| *pyrH* | HSISS4_00354 | Uridine monophosphate kinase | -0.94 | -3.03 |
| *sipA* | HSISS4_01673 | Secretory signal peptidase | -0.87 | -2.63 |
|  | HSISS4_00898 | Permease | 0.05 | -2.73 |
|  | HSISS4_00523 | Hypothetical protein | 0.16 | -3.82 |
|  | HSISS4_00888 | Hypothetical protein | -0.03 | -3.30 |

<sup>a</sup> Fitness-cost scores were computed with the MAGeCK algorithm (41) by comparing the total depletion of gRNAs per gene in the mock condition with the library-induced condition (Ci).
<sup>b</sup> Standardized (Std) residuals (cut-off value < -2.5) were calculated as the deviation from the linear regression performed with the fitness-cost scores for conditions with library induction (Ci) and with library induction together with competence induction (Ci+C).

### Screening based on β-galactosidase activity

The last screening strategy was based on β-galactosidase (β-Gal) activity that allows the colorimetric evaluation of competence activation level in individual clones on plates with low selective pressure. For this purpose, a strain was generated by plugging the promoter of *comX* in front of the native *lacZ* gene (P*_comX_-lacZ*) together with a P*_comX_* luciferase reporter system (P*_comX_-luxAB*) into the dCas9 and xylose-inducible competence strain (Fig. 1C). We transformed the gRNA library into this genetic background and spread it onto M17GL plates supplemented with IPTG, xylose, and X-gal for detection of β- Gal activity. We examined ∼94,000 isolated colonies, searching for white and dark blue phenotypes. While white phenotype is associated to targeted genes required for competence activation, dark blue phenotype is related to targeted genes repressing competence development. We next re-isolated the selected colonies to confirm their phenotypes and ended up with 141 dark blue and 68 white clones. We sequenced their gRNAs to identify the interference target and quantified their inhibition effects on P*_comX_* activation thanks to the P*_comX_*-*luxAB* module present in the strain. To this aim, we slightly overexpressed *comR* with xylose thanks to the P*_xyl2_-comR* module and induced the gRNA-based inhibition system by adding IPTG. We compared the specific luciferase activity of all the selected clones to the initial strain harboring no gRNA. The sequence of the gRNAs, their identified targets, and their fold-changes in P*_comX_* activation are displayed in Table S5.

Since both frequency of selected gRNAs targeting the same gene and fold-change in P*_comX_* activation were relevant features to identify new competence regulators, we combined those two parameters in the same analysis. On one hand, we calculated the mean fold-change in P*_comX_* activation for all gRNAs targeting the same gene. On the other hand, we counted the number of selected gRNAs targeting the same gene. In addition, we normalized the count by the total expected number of gRNAs of the library targeting each defined gene to avoid any gene-size bias (higher probability to encounter a gRNA targeting a high-size gene) (Table S6). We plotted those two variables (activation fold-change vs normalized gRNA frequency) and validated the screen by finding most of the proximal effectors of the ComRS system, i.e. *comR*, *amiACDEF, clpC* and P*_comX_* (Fig. 2B and Table S6) (7, 16, 42). We next applied an arbitrary cut-off (normalized count > 0.02 and log_2_(FC) > 0.5) to find the most significant genes with an antagonist function towards competence (Table 3). We thereby selected several players whose role in competence inhibition was also suggested with the gRNA depletion screen, such as the phosphate transporter system (*pstC1*) and the serine-threonine kinase (*stkP*) genes. Strikingly, the mannose/glucosamine PTS transporter operon (*manLMN*) was particularly overrepresented, even though absent from the two previous screens. This result might be an artefactual consequence of a carbon metabolism shift enhancing xylose uptake ultimately resulting in higher *comR* induction, but could also be due to a link between mannose catabolism and competence as reported in *S. mutans* (30).

**Table 3.**
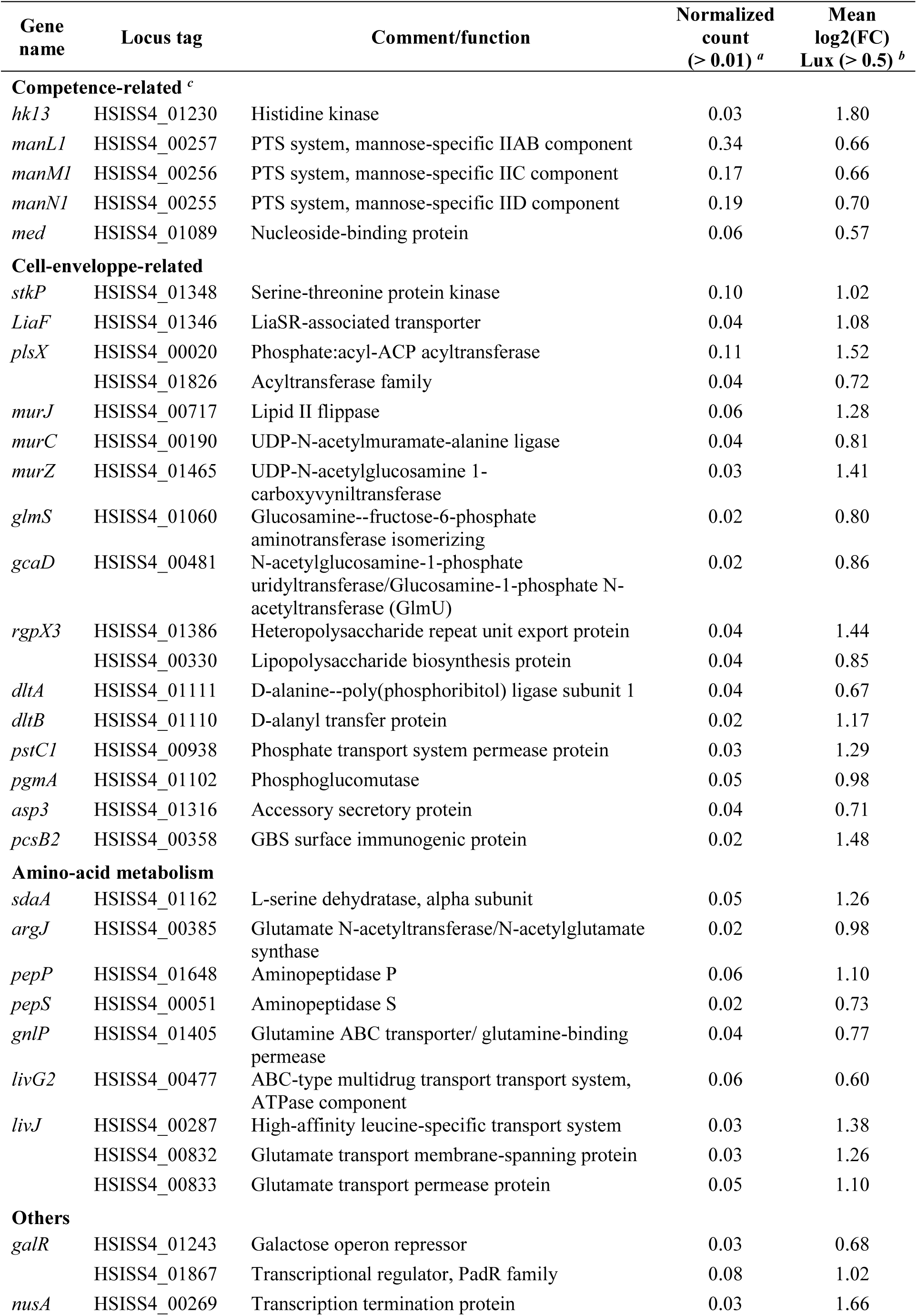
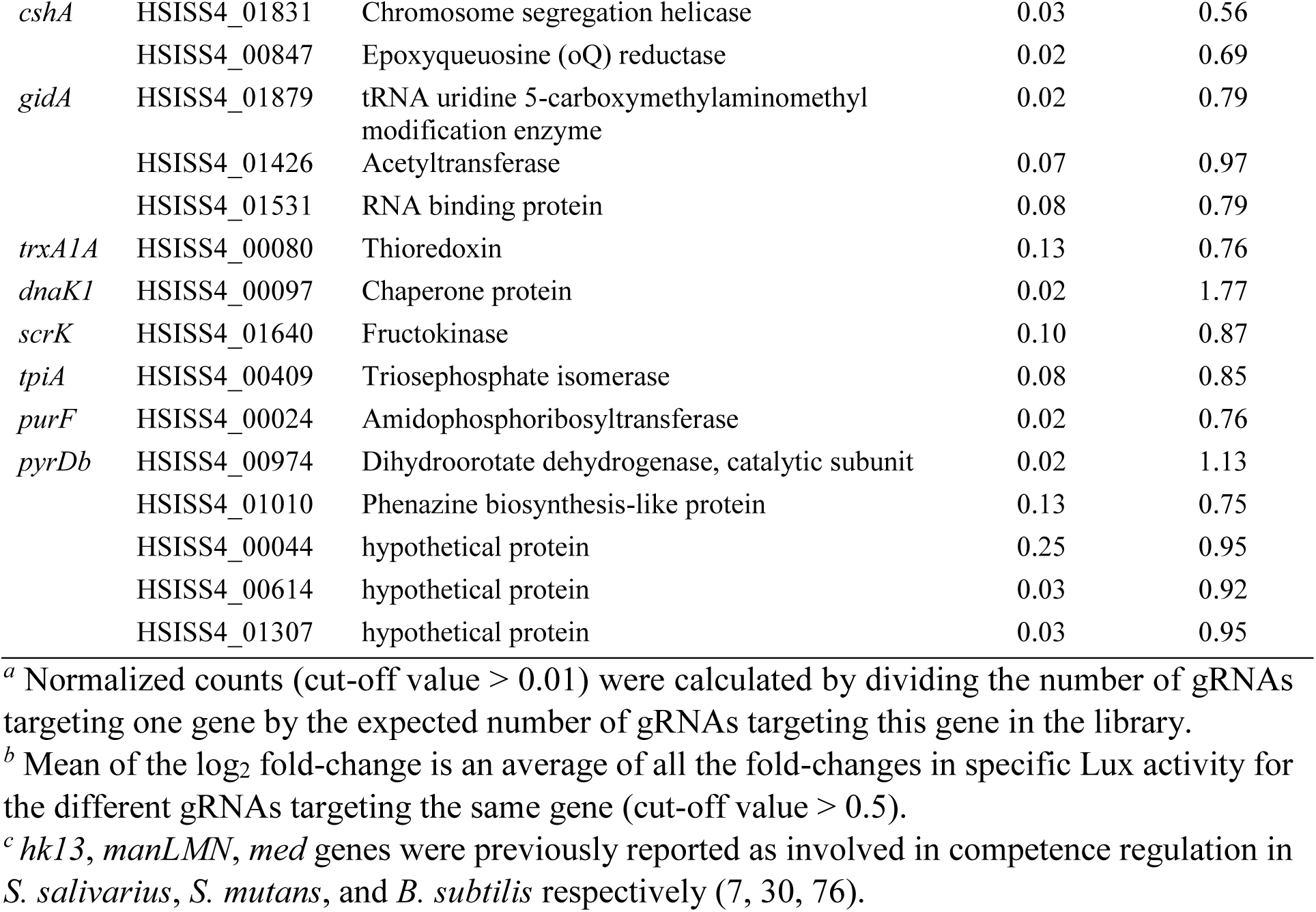
Identification of competence-associated antagonist genes from β-Gal screen.

### Cell-wall integrity is a signal for competence

Since many genes were identified to affect competence by the three screening approaches, we used the Clusters of Orthologous Genes (COG) database (44) to classify them by general function. For this analysis, we selected only the genes whose inhibition is expected to induce competence (i.e. all the genes from the transformation screen, genes with standardized residuals < -2.5 from the growth screen, and genes with log_2_(FC) > 0.5 and normalized count > 0.02 from the colorimetric screen). We next assessed the importance of the different pathways for competence activation. For this purpose, we counted the number of genes per screen involved in one COG function and normalized it by the total number of genes within this COG function in the HSISS4 genome (Fig. 3A). This analysis indicated that the most represented function was cell wall/membrane/envelope biogenesis (23% of all the genes highlighted vs ∼5% at the whole genome level). Furthermore, we observed that overlapping genes between gRNA depletion and β-Gal screens were all directly or indirectly involved in cell-envelope assembly. Indeed, we identified in both screens the *dltA* and *dltB* genes involved in teichoic acid D-alanylation (45), the phosphoglucomutase *pgmA* gene involved in the biosynthesis of extracellular polysaccharides (46), the cell wall-related serine-threonine kinase *stkP* gene (47, 48), and the phosphate transporter *pstC1* gene with an important role for poly(glycerophosphate) teichoic acid synthesis (49) (Fig. 3B).

**Figure 3.**
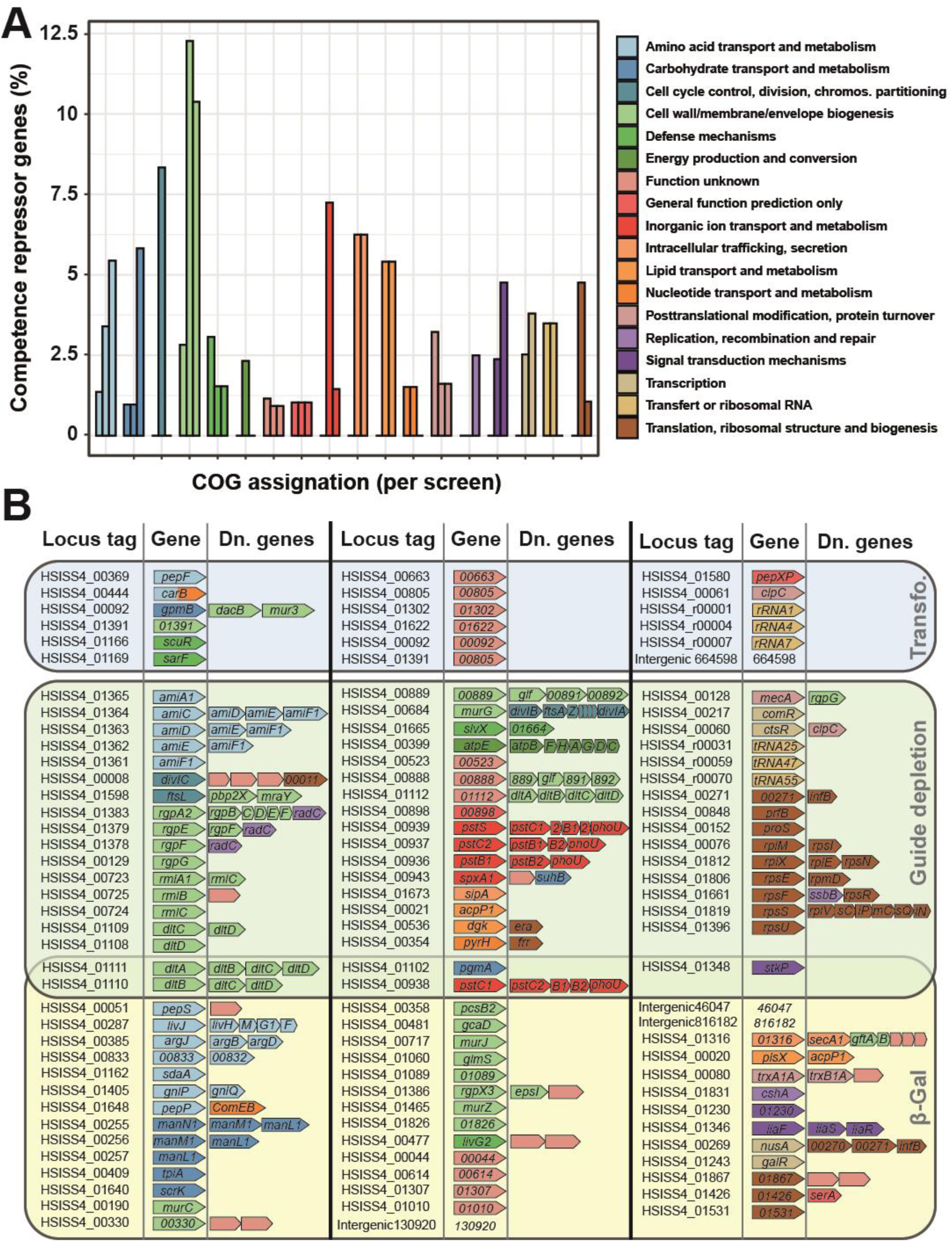
Functional assignation of competence repressor genes from CRISPRi screens. **A.** Relative abundance of COG-assigned genes. A COG assignation was associated to every gene from the HSISS4 genome. For each COG type, the proportion (%) of selected genes with a defined screen was calculated against all the genes with this COG assignation of the genome. This proportion is displayed per screen (1^st^ bar, transformation screen; 2^nd^ bar, gRNA depletion screen; 3^rd^ bar, β-Gal screen). **B.** Details of all selected genes displayed per screen. Operons (Dn. genes) are shown since CRISPRi also silences downstream genes. Genes are colored according to their COG assignation.

We next drew a more precise view of the different cell wall pathways targeted by gRNA that presumably lead to competence activation. We found that genes involved in the synthesis of peptidoglycan, teichoic acids, and extracellular polysaccharides were all affected (Fig. 4). In parallel, key sensors (StkP, LiaFSR) or mediators (SpxA1) known to be triggered by cell wall alterations were also identified in the screens (24, 50–54), suggesting a possible link between cell wall integrity and ComRS activation.

**Figure 4.**
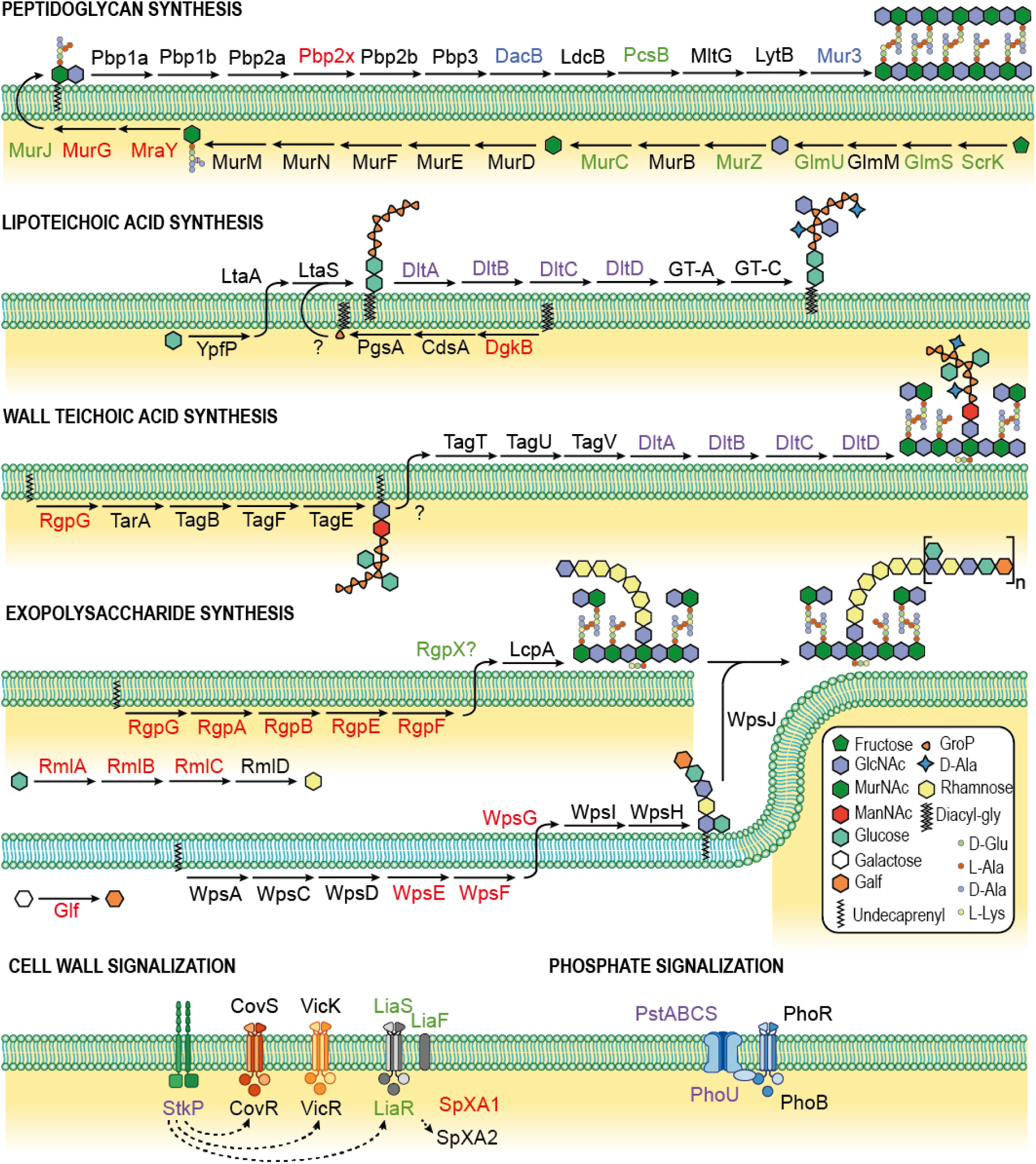
Cell-wall pathways and competence negative modulators from CRISPRi screens. Major cell-wall biosynthesis and signalization pathways are depicted. Proteins selected by the transformation, gRNA depletion, and β-gal screens are shown in blue, red, and green, respectively. Proteins selected in both gRNA depletion and β-gal screens are shown in light violet. In the absence of literature for complete reconstructed pathways, lipoteichoic acid synthesis is based on knowledge from *Staphylococcus aureus,* wall teichoic acid from *B. subtilis* 168 and polysaccharide synthesis from *L. lactis*. HSISS4_00889, 00890, 00891 and 00892 were renamed with *Lactococcus lactis* homologs WpsG, Glf, WpsE and WpsF, respectively. GlcNAc, *N*-acetylglucosamine; MurNAc, *N*-acetyl muramic acid; ManNAc, *N*- acetylmannosamine; Galf, Galactofuranose; Diacyl-gly, Diacyl-glycerol; GroP, Glycerol- phosphate.

### StkP controls *comX* expression through muropeptide binding

Since StkP was highlighted in two screens with multiple different gRNAs and is cell wall-associated, we decided to further investigate its link with competence activation. Serine-threonine kinases are pleiotropic regulators that control key cellular processes such as dormancy, virulence, cell division, and cell wall synthesis through protein phosphorylation (47, 48). In *S. salivarius*, only one serine-threonine kinase homolog is present and displays PASTA motifs shown to bind muropeptides in *Bacillus subtilis* (55). Besides, StkP of *S. pneumoniae* has been shown to coordinate cell wall synthesis and septation (24, 56–58) while an unclear link with competence has been suggested in *S. mutans* and *S. pneumoniae* (19, 23, 24).

In a first set of experiments, we transferred a gRNA targeting *stkP* in a strain harboring the dCas9 module (P_F6_-*lacI* P*_lac_-dcas9*) together with a luciferase reporter of the transcriptional activity of *comR* (P*_comR_-luxAB*) or *comX* (P*_comX_-luxAB*) and the xylose-inducible module allowing competence activation (P*_xyl2_-comR*). Monitoring activation of those two promoters with or without *stkP* inhibition suggested that StkP level influences *comX* expression but has no impact on *comR* transcription (Fig. 5A). We next used the same *comX* reporter strain with increasing xylose concentrations for *comR* induction and measured P*_comX_* activation with or without *stkP* inhibition (Fig. 5B). Stronger inhibitions of *stkP* were recorded for lower ComR levels, suggesting that ComR overproduction interferes with the StkP-mediated regulation of *comX*.

**Figure 5.**
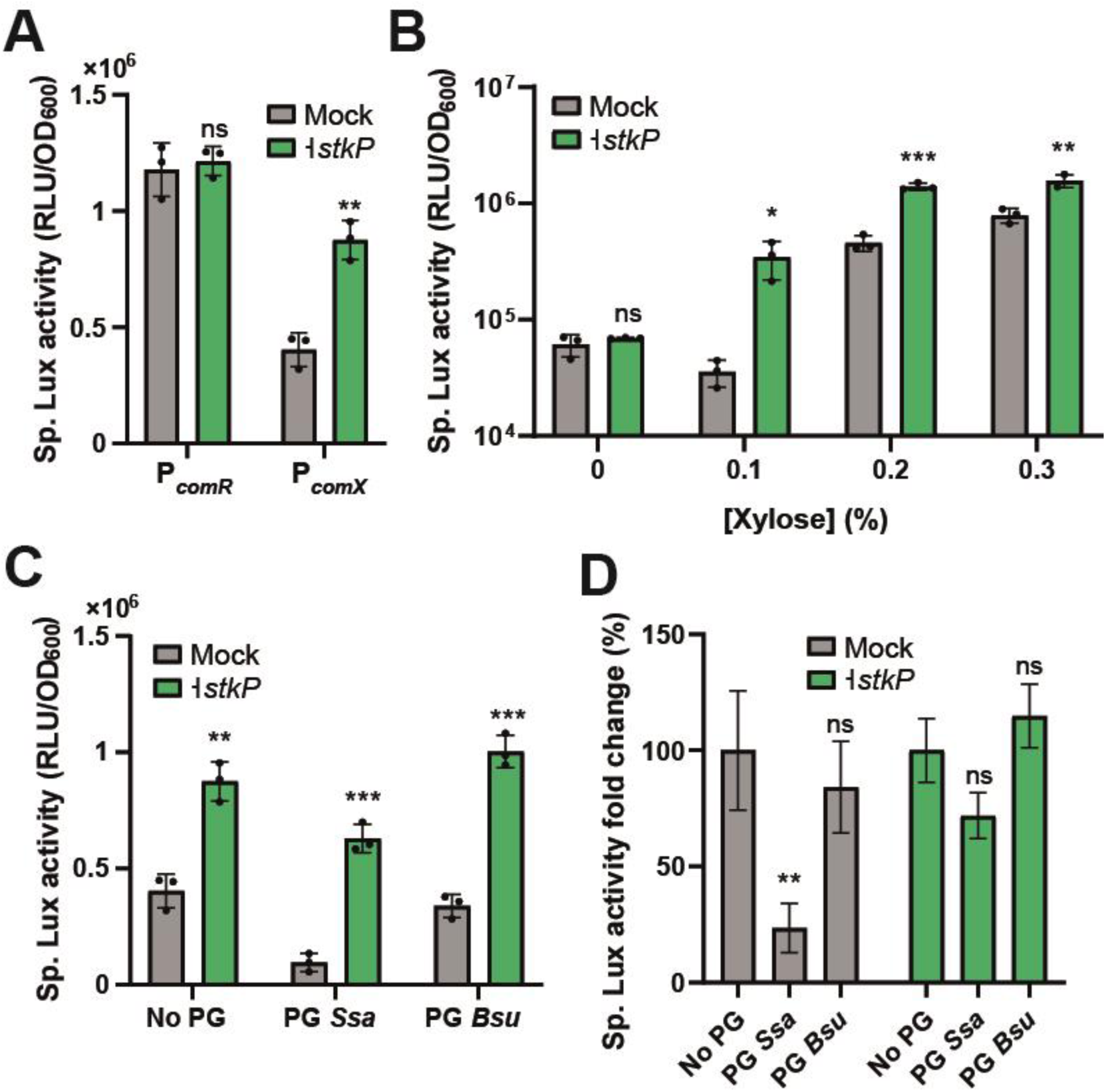
StkP controls *comX* expression by sensing peptidoglycan extracts. **A.** Effect of *stkP* inhibition on *comR* and *comX* expression. A dCas9 module (P_F6_*-lacI*; P*_lac_-dcas9*) together with a gRNA targeting *stkP* (P_3_-*g_23*) were used to inhibit *stkP* transcription. The dCas9 interference system was associated to a P*_comR_*-*luxAB* or a P*_comX_-luxAB* reporter fusion together with a xylose-inducible *comR* module (P*_xyl2_-comR*). Mock denotes the same strain without dCas9 interference. **B.** Effect of ComR level on StkP-mediated activation of *comX*. The P*_comX_- luxAB* P*_xyl2_-comR* strain (described in A) was incubated with various xylose concentrations (0, 0.1, 0.2, 0.3%) with or without *stkP* inhibition. **C.** Effect of peptidoglycan (PG) extracts on StkP-mediated activation of *comX*. PG extracts of *S. salivarius* (PG *Ssa*) or *B. subtilis* (PG *Bsu*) were added to a culture of the P*_comX_-luxAB* strain (described in A) at a final concentration of 0.3 mg/ml. **D.** Specific Lux activity (%) calculated between the culture with no addition of PG extracts (No PG, 100%) and the related condition. Percentages were calculated with the data presented in C. For P*_comX_-luxAB* activation, xylose 0.25% was used except if else stated. For CRISPRi *stkP* inhibition, IPTG 1 mM was used. Dots denote technical triplicate values for mock and biological triplicate values for mutants, ± standard deviation. Statistical *t*-test was performed for each condition in comparison to related control (Ctrl: mock; panels A, B, and D) or one-way ANOVA with Dunett’s test for multiple comparison (Ctrl: No PG; panel E). ns, non-significative; *P* > 0.05; *, *P* < 0.05; **, *P* < 0.01; ***, *P* < 0.001.

Since StkP was shown to bind specific muropeptides via its PASTA domains (55), we investigated if the addition of peptidoglycan extracts was able to modulate competence. To this aim, we purified peptidoglycan from *S. salivarius* (L-Lys pentapeptide) or *B. subtilis* (Meso- DAP pentapeptide) and measured the activation of the P*_comX_-luxAB* reporter strain with (-| *stkP*) or without (mock) dCas9 interference on *stkP* expression (Fig. 5C). While *S. salivarius* peptidoglycan could decrease P*_comX_* activation, *B. subtilis* extracts (negative control) did not result in a significative reduction. Moreover, adding peptidoglycan from *S. salivarius* prevented P*_comX_* repression when *stkP* was inhibited, suggesting that StkP mediates the signalization (Fig. 5D).

Altogether, these results suggest that StkP interferes with the transcriptional activity of the ComR**·**XIP complex by an unknown mechanism, which is modulated by the binding of specific muropeptides.

## DISCUSSION

How QS modules integrate multiple inputs to fine-tune their sensitivity and optimize collective behavior is a challenging topic. In this work, we performed a genome-wide screen coupled to three different readouts to uncover key triggers of ComRS-mediated competence activation. Using a conditional mutant library, we highlighted a connection between cell wall biogenesis and competence activation. Moreover, we uncovered a link between muropeptide sensing via the serine threonine StkP and competence development. Those pieces of evidence suggest a key role of cell wall stress on the competence response (Fig. 6).

**Figure 6.**
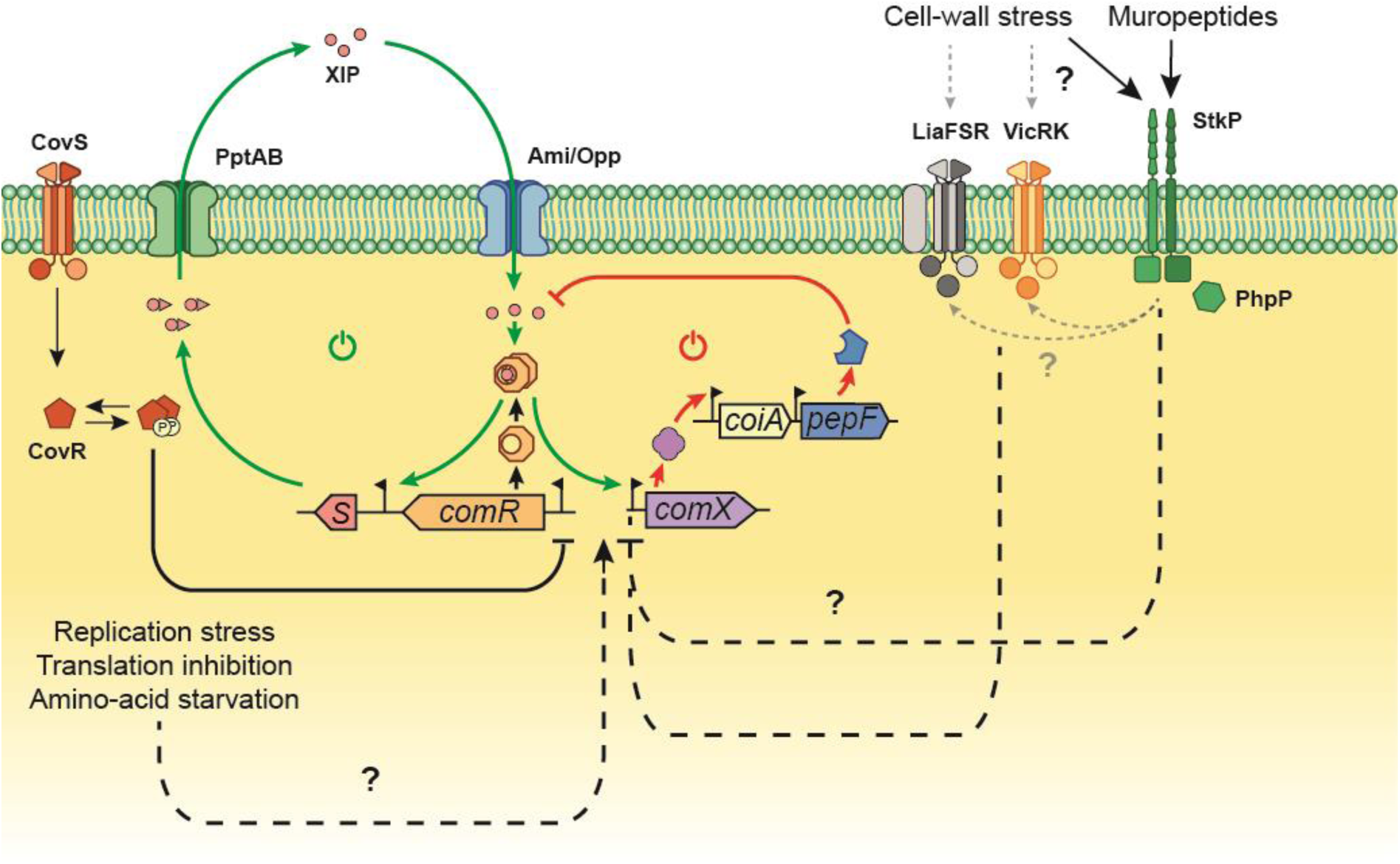
Model of competence regulation integrating cell-wall sensors and physiological stresses in *S. salivarius*. Upon CovRS repression release, ComR reaches a threshold concentration allowing the activation of a positive feedback loop (green arrows, power-on icon). The positive loop is triggered by XIP binding to ComR, producing the ComR•XIP complex which activates *comS* transcription. ComS is then exported by the transporter PptAB and maturated. The mature XIP pheromone can then enter the cell by the oligopeptide generic transporter Ami/Opp and bind ComR to enhance the loop. In parallel, the ComR•XIP complex will trigger the transcription of *comX*, encoding the central regulator of competence. This will activate all the late genes necessary for natural transformation including the *coiA-pepF* operon. PepF accumulation will result in XIP degradation, generating a negative feedback-loop (red arrows, power-off icon) on the ComRS system, ultimately leading to competence exit. In parallel, cell-wall stress and/or free muropeptides can be sensed by the serine-threonine kinase StkP, LiaFSR and VicRK to modulate the transcriptional activation of *comX*, most probably via interfering with the activity of the ComR•XIP complex. Other physiological stresses such as replication stress, translation inhibition or amino-acid starvation were also identified as conditions that could activate competence development.

To discover novel players involved in competence regulation, we built a CRISPRi-based library and performed three types of screen in parallel. The interference technology offers several advantages over the classical random transposon mutagenesis (59) but the primary one is the production of conditional mutants allowing the study of essential/deleterious genes. Considering the transformation screen, the library was transiently induced, dampening the toxic-acquired phenotype due to constitutive activation of natural transformation. This strategy provided a direct screening method for DNA integration and allowed us to select gRNAs targeting essential genes among which *pepF*, a gene essential for competence shut-off recently discovered in *S. salivarius* (40). In addition, we also selected two different gRNAs targeting *clpC,* a gene encoding a component of the MecA-ClpCP machinery responsible for ComX degradation (39, 60). Those results confirm the roles of PepF and ClpC to prevent spontaneous competence activation at the early and late stage of competence, respectively (40, 42, 60). Moreover, novel competence modulators were identified such as a putative bactoprenol glucosyltransferase and 3 hypothetical proteins. Specifically, interference on the putative bactoprenol glucosyltransferase resulted in a high transformation rate (∼10^-2^, Table 1), suggesting an important role of this player for competence control. Although the transformation screen displays interesting features to select essential genes connected to competence development, it would require a massive number of cells to ensure a complete coverage of the high-density gRNA library. This issue is not present in the gRNA depletion screen, where high-throughput NGS is exploited to map and quantify all the gRNAs, generating a complete picture at the genome scale. Nevertheless, the identification of the genes is based on the competence-related toxic phenotype. This feature could limit the detection of essential genes whose inhibition has a high fitness cost. Of note, the competence-associated toxicity used in the gRNA depletion screen could explain some intriguing results. While NGS data showed a depletion of gRNAs targeting genes involved in the downregulation of competence such as *mecA*, the depletion of gRNAs targeting crucial genes for competence activation (e.g. *comR* or the *ami/opp* operon) was counterintuitive. To reconcile these findings, we reasoned that a lack of functional competence goes along with an impairment in bacteriocin immunity. Consequently, the gRNA depletion will also include bacteriocin/immunity loci and key players required for competence activation (Table 2 and Table S3). Finally, as the colorimetric β-gal test is based on P*_comX_* activity and visual selection, this screen drastically reduces any fitness bias. To sum up, this work highlights the added value of combining different screening approaches to unveil the largest set of candidate genes connected to competence control.

The three screens converge to select gRNAs involved in key envelope biogenesis processes and its control by cell wall sensors (Fig. 4). The connection between cell wall and competence has only been reported in a similar experiment with Tn-seq in *S. mutans* (61). However, the authors report that inactivation by transposon insertion of the cell wall related genes *pknB* (homolog of *stkP*), *rgpL*, *dltA* and *liaS* results in a lower activation of competence, contrasting with the results obtained in this study. Opposite effects of competence regulators in *S. salivarius* and *S. mutans* have already been reported for the CovRS system (7, 62) and showcase that species have evolved control mechanisms in line with their own lifestyles. Aside from the cell wall synthesis, several other pathways were highlighted (Fig. 6). One of them is translation, with several important players targeted (rRNAs, tRNAs, peptide chain release factor, ribosomal proteins, and tRNA synthetases). This correlation is interesting in the light of the work of Stevens and coworkers, who showed that translation fidelity impairment promotes competence activation in *S. pneumoniae* (63). In addition, important genes involved in chromosome replication/segregation (*priA*, *cshA/rarA*, *scpB*) and DNA repair (*mutL*, *mutT*, *dinP*) were also underlined by the screens (Table 3, Tables S4 and S6). Replication stress was previously shown to induce pneumococcal competence, but the exact mechanism remains unclear and involves *comCDE* gene dosage control and/or a role for arrested and unrepaired replication forks (64, 65). The screens did highlight a role for enzymes or transporters involved in amino acid biosynthesis or uptake for arginine (CarB and ArgJ), glutamine (GlnP), glutamate (HSISS4_00833 and 00832) and leucine (LivJ). Amino acid starvation is known to trigger the stringent response via RelA and the production of (p)ppGpp alarmones (66), which was shown to influence competence regulation in *S. mutans* (67). Altogether, the screens performed here suggest that *S. salivarius* competence control relies on the sensing of various alterations of key metabolic/physiological functions, reinforcing the view that competence activation could be seen as a general stress response in streptococci.

In this work, we specifically investigate StkP, a key sensor of cell wall integrity in *S. pneumoniae* (54). In streptococci, StkP was also shown to phosphorylate classical response regulators of two-component systems such as the virulence regulator CovR in *Streptococcus agalactiae* and *Streptococcus pyogenes* (68, 69), the cell wall regulator VicR in *S. mutans* and *S. pneumoniae* (24, 54), and the competence regulator ComE in *S. pneumoniae* for which StkP phosphorylation triggers a distinct regulon from the aspartate phospho-transfer mediated by ComD (22). The pleiotropic effects of StkP and its involvement in major cellular processes is probably the reason why its impact on competence has been reported but remains controversial in *S. pneumoniae* (19, 22, 23). In *S. salivarius*, we showed that *stkP* depletion promotes a higher *comX* activation without major effect on *comR* expression. This suggests a mechanism acting directly on the ComR sensor by increasing its trans-activator properties. This hypothesis is strengthened by the fact that ComR overexpression curtails the effect of StkP on *comX* activation (Fig. 5B). The exact process remains to be discovered, even if it suggests a direct effect of StkP on ComR. Two non-exclusive mechanisms could explain the control of competence by StkP in *S. salivarius*. On one hand, the kinase could sense directly or indirectly a disfunction in the cell wall synthesis. Beside a direct effect on ComR, this impairment could also be transmitted to other cell wall sensors. Interestingly, two of these sensor systems (i.e. VicRK and LiaSRF) were highlighted in the β-gal screen (Tables 3 and S5) and were previously shown to affect competence activation in *S. salivarius* (Fig. 6) (7). On the other hand, the kinase could also control competence as a muropeptide signaling system. Our experiments with peptidoglycan extracts (Fig. 5C and D) advocate for this possibility as a high concentration of self-muropeptides inhibits competence in a StkP-dependent manner. In line with this, competence in streptococci is transiently activated during the early exponential growth but could not be triggered in stationary phase when the extracellular muropeptide concentration is expected to be high (14, 16, 70). This may suggest that StkP acts as a growth phase sensor to extinct competence at high cell density (Fig. 6).

To conclude, we showed the large potential of combining a genome-wide CRISPRi strategy with multiple screening approaches to connect essential genes involved in physiological pathways to competence development. Besides the well-established ComRS-ComX regulatory pathway, we revealed that disturbance of general functions such as cell envelope assembly, amino-acid metabolism, translation, and replication modulates competence activation in *S. salivarius*. This work strengthens our view that competence is a general adaptative response that ensures survival in a broad range of stress conditions. Moreover, the identification of a large set of “competence-associated” genes paves the way to understand novel regulatory cascades interconnecting cell-proliferation impairment and competence activation such as illustrated here for the role of the serine/threonine kinase StkP in cell wall-mediated competence modulation.

## MATERIALS AND METHODS

### Bacterial strains, plasmids, sgRNA and oligonucleotides

Bacterial strains, plasmids, sgRNA and oligonucleotides used in this study are listed and described in supplemental material (Tables S7A-D).

### Growth conditions and competence induction

*S. salivarius* HSISS4 (71) and derivatives were grown at 37°C without shaking in M17 (Difco Laboratories, Detroit, MI) or in chemically-defined medium (CDM) (72) supplemented with 1% (w/v) glucose (M17G, CDMG, respectively). Chromosomal genetic constructions were introduced in *S. salivarius* via natural transformation (73). We added D-xylose (0.1 to 1%; w/v), IPTG (1 mM), spectinomycin (200 µg/ml), chloramphenicol (5 µg/ml) or erythromycin (10 µg/ml), as required. The synthetic peptides (purity of 95%), were supplied by Peptide 2.0 Inc. (Chantilly, VA) and resuspended first in dimethyl-formamide (DMF) and diluted in water to reach low DMF concentration (final of 0.02%). Solid plates inoculated with streptococci cells were incubated anaerobically (BBL GasPak systems, Becton Dickinson, Franklin lakes, NJ) at 37°C.

To induce competence, overnight CDMG precultures were diluted at a final OD_600_ of 0.05 in 500 µl of fresh CDMG and incubated 100 min at 37°C. Then, the pheromone sXIP (LPYFAGCL) and DNA (Gibson assembled PCR products or plasmids) were added and cells were incubated for 3 h at 37°C before plating on M17G agar supplemented with antibiotics when required.

### Library design and construction

The gRNA library was designed by selecting all the 20 nt followed by a PAM sequence within the genome of HSISS4. Since it was shown that efficient interference in CDSs occurs only with gRNAs targeting the coding strand (34), we filtered the library to keep only gRNAs targeting the coding strand in CDSs and targeting both strands in intergenic regions. We ended up with a high-density library of 83,104 gRNAs resulting in a theoretical base pairing every 25 pb on the HSISS4 genome. Of note, we chose to use a high-density library to target unknown small genetic elements such as siRNA or small peptides. We ordered the gRNAs as single strand DNA (Twist BioSciences) and amplified the oligo-pool thanks to common upstream and downstream region using primers AK475 and AK476. To reduce any amplification sequence-bias, we used 10 cycles of amplification.

The PCR products were then purified (Monarch kit, NEB) and Gibson-joined to the pre-amplified upstream homologous region of the neutral locus *gor* (downstream of *HSISS4_00325*) containing an erythromycin resistance gene and to the downstream homologous region of the same locus fused to a P_3_ constitutive promoter. We performed 20 independent Gibson assemblies, which were latter transformed by natural transformation into 20 independent cultures of HSISS4 strains containing at least a lactose-inducible dCas9 module (P_F6_-*lacI*; P*_lac_-dcas9*) (7). Supplemental genetic constructions (P*_xyl2_-comR*, P*_comX_-lacZ*, P*_comX_- luxAB*) were present in those strains depending on the screening strategy. For every library produced, we calculated the number of transformants to obtain at least 15-fold transformants over the diversity rate, ensuring theoretically that 99,9% of the diversity would be present in the library (74). We finally pooled all the transformants in Phosphate Buffered Saline (PBS, 137 mM NaCl, 2.7 mM KCl, 10 mM Na_2_HPO_4_, 1.8 mM KH_2_PO_4_) and measured OD_600_ prior to storage at -80°C.

### CRISPRi transformation screen

For the spontaneous transformation screen, we first introduced the gRNA library in the lactose-inducible strain harboring a P*_comX_* luciferase reporter system (P_F6_*-lacI,* P*_lac_-dcas9*, P*_comX_-luxAB*). We next diluted the cells in 15 ml of fresh CDMG supplemented with IPTG 1 mM at an OD_600_ of 0.01. We grew this culture at 37°C for 8 h and added every 30 min a PCR-amplified product consisting of a chloramphenicol cassette with 2000-pb up and down homologous recombination arms at a final concentration of 0.25 nM. We centrifugated this culture, plated it on chloramphenicol plates and grew it overnight. Colonies were picked and donor DNA integration was confirmed by PCR. We next amplified the locus containing the gRNAs before Sanger sequencing.

### CRISPRi gRNA depletion screen

For the gRNA depletion analysis, we used the same strain as described in the previous section. After introducing the gRNAs in this background, we spread the resulting library onto three different solid media (M17G; M17G IPTG 100 µM; M17G IPTG 100 µM xylose 1%) resulting in an average of 9.6 × 10^6^ CFU per large plate. Technical replicates (*n* = 4) were incubated 16 h at 37°C to yield an estimated mean of ∼12 generations. Cells were then collected, pooled in PBS buffer and homogenized for each replicate. After genomic extraction (GenElute, Sigma Aldrich) from at least 1.5 × 10^9^ CFU per replicate, we PCR-amplified the locus containing the gRNAs prior to their deep sequencing. We used an optimized PCR protocol with high primer concentration (5 µM), low template genomic DNA (2 ng/µl), and low number of cycles (15 cycles) to avoid any chimeric products due to the highly randomized gRNA sequences. The 219-pb amplicons were next gel-purified (Monarch DNA Gel Extraction kit, NEB) and sent with a minimum amount of 4 pmol for Illumina sequencing (GeneWiz). High-Seq Illumina sequencing was performed with 30% Phix and generated an average of 30 M reads per replicate. All gRNA sequencing data was deposited in the GEO database under accession number GSE204976.

### CRISPRi β-galactosidase activity screen

We first produced a new genetic background by introducing into the strain described in the previous section an ectopic copy of *comR* under the control of a xylose-inducible promoter (P*_xyl2_-comR*) together with a chloramphenicol resistance cassette at the neutral locus *suc* (upstream of *HSISS4_01641*). We next fused the promoter of *comX* to the native *lacZ* gene (P*_comX_-lacZ*) together with a spectinomycin resistance cassette and introduced the gRNA library into this strain. The resulting library was spread on M17 0.5% glucose, 0.5% lactose (M17GL), IPTG 100 µM, xylose 1%, and X-gal 100 mg/ml for screening dark-blue (highly competent) and white (competence loss) colonies. A total of 158 dark blue and 155 white clones from the screening of ∼94,000 colonies were re-isolated for phenotype confirmation. Luciferase tests (P*_comX_-luxAB*) were performed in comparison with the parental strain harboring no gRNA. Clones with the most dissimilar luciferase phenotypes (141 dark blue and 68 white clones) were selected and gRNAs were amplified by PCR for Sanger-sequencing.

### NGS analysis

We used the MAGeCK algorithm to map the reads on the HSISS4 genome (41). Approximatively 30% of total reads were mapped, producing about 10 M reads per replicate. Following the MAGeCK guidelines, we next pooled the reads from the 4 replicates, ultimately generating a total of 40 M reads per condition. In a first analysis, we compared the gRNA depletion in the IPTG-induced condition with the mock to determine all the essential genes from strain HSISS4. For the sake of clarity, we only compared gRNAs targeting CDSs, since gRNAs targeting intergenic regions are much more complicated to determine. We next compared the depletion of gRNAs for each gene in the IPTG- and IPTG-xylose-induced conditions to the mock condition thanks to the MAGeCK algorithm. The depletion scores generated per gene for the two induced conditions were then plotted against each other and a linear regression was fitted to the plot (*lm* function, R package) and outliers were identified by standardizing the residuals.

### COG analysis

The conserved domain database of NCBI was used to infer functions of the genes from the genome of HSISS4 (44, 75) and only the highest scored function for each gene were retained. The number of genes of the whole genome involved in each function prediction was then calculated, generating a function prediction frequency matrix. This matrix was then used to weight the number of genes with a specific predicted function highlighted in the different screens.

### Luciferase assay

Overnight precultures were diluted at a final OD_600_ of 0.05. A volume of 300 µl of culture was transferred in the wells of a sterile covered white microplate with a transparent bottom (Greiner, Alphen a/d Rijn, The Netherlands). These culture samples were supplemented with D-xylose, IPTG or peptidoglycan extracts if stated. Growth (OD_600_) and luciferase (Lux) activity (expressed in relative light units, RLU) were monitored at 10 min intervals during 8 to 24 h in a Hidex Sense microplate reader (Hidex, Lemminkäisenkatu, Finland). Specific Lux activity were obtained by dividing Lux activity by the OD_600_ and summing all the data obtained over time. When stated, biological or technical triplicates were averaged. Statistical analyses of simple and multiple comparisons to the control mean were performed with *t*-test (unilateral distribution, heteroscedastic) and one-way ANOVA with Dunnett’s test, respectively. For both, standard deviations and *P* values were calculated.

### Transformation test

The CDMG preculture of HSISS4 and derivatives was diluted in 500 µl of CDMG supplemented with 1 mM IPTG at an OD_600_ of 0.005. The culture was grown at 37°C for 8 h and we added every 30 min a PCR-amplified product consisting of a chloramphenicol resistance cassette surrounded by up and down homologous recombination arms (2000 pb each) at a final concentration of 0.25 nM. We next performed serial dilution of the culture and spread the various dilutions on M17G plates supplemented with or without chloramphenicol 5 µg/ml. We next calculated the transformation rate based on the CFU numbers of the two plates.

### Peptidoglycan extracts

Peptidoglycan extracts were prepared as previously reported (55). Cultures of 100 ml of *S. salivarius* HSISS4 or *B. subtilis* 168 were grown to an OD_600_ of ∼1.2 in M17 or LB media, respectively. Cells were collected by centrifugation, washed with 0.8% NaCl, resuspended in hot 4% SDS, boiled for 30 min, and incubated at room temperature overnight. The suspension was then boiled for 10 min and the SDS-insoluble cell-wall material was collected by centrifugation at 12,000 *g* for 15 min at room temperature. The pellet-containing cell wall peptidoglycan was washed four times with water and finally resuspended in 1 ml sterile water. The resuspended peptidoglycan was next digested with mutanolysin (10 μg/ml) overnight at 37°C prior to inactivation of mutanolysin at 80°C for 20 min.

## ACKNOWLEDGEMENTS

The work of PH was supported by the Belgian National Fund for Scientific Research (FNRS, grant PDR T.0110.18) and the Concerted Research Actions (ARC, grant 17/22-084) from Federation Wallonia-Brussels. AK held a doctoral fellowship from FNRS (FRIA fellowship). PH is Research Director at FNRS. The funders had no role in study design, data collection and interpretation, or the decision to submit the work for publication.

We thank Marcello Mora for its advices on non-specific DNA amplification.

## AUTHOR CONTRIBUTIONS

AK and PH conceived and designed the study. AK, AW, ML, BD, and JM carried the laboratory work. AK, JWV, JM and PH analyzed and interpreted the data. AK, JWV, JM and PH wrote and revised the manuscript. All authors read and approved the final manuscript.

## SUPPLEMENTAL MATERIAL

### Supplementary Figures

**Figure S1. Random chromosomal distribution of the gRNA library**

**Figure S2. Gene-associated gRNA depletion scores**

**Figure S3. Linear regression of gene-associated gRNA depletion scores**

### Supplementary Tables

**Table S1. List of oligonucleotides used for the genome-wide CRISPRi strategy**

(separate .xls file)

**Table S2. List of gene-associated gRNA depletion scores from library activation**

(separate .xls file)

**Table S3. List of gene-associated gRNA depletion scores from library and competence activation**

(separate .xls file)

**Table S4. List of competence-associated genes (standardized residuals) from the gRNA depletion screen**

(separate .xls file)

**Table S5. List of gRNA and their targeted genes from the β-Gal screen**

(separate .xls file)

**Table S6. List of normalized competence-associated genes from the β-Gal screen**

(separate .xls file)

**Table S7. List of bacterial strains (A), plasmids (B), oligonucleotides (C), and PCR fragments (D) used in this study**

## REFERENCES

1. Huang R, Li M, Gregory RL. 2011. Bacterial interactions in dental biofilm. Virulence 2:435–444.

2. Huttenhower C, Gevers D, Knight R, Abubucker S, Badger JH, Chinwalla AT, Creasy HH, et al. 2012. Structure, function and diversity of the healthy human microbiome. Nature 486:207–214.

3. Kommineni S, Bretl DJ, Lam V, Chakraborty R, Hayward M, Simpson P, Cao Y, Bousounis P, Kristich CJ, Salzman NH. 2015. Bacteriocin production augments niche competition by enterococci in the mammalian gastrointestinal tract. Nature 526:719–722.

4. Bettenworth V, Steinfeld B, Duin H, Petersen K, Streit WR, Bischofs I, Becker A. 2019. Phenotypic Heterogeneity in Bacterial Quorum Sensing Systems. J Mol Biol 431:4530–4546.

5. Doganer BA, Yan LKQ, Youk H. 2016. Autocrine Signaling and Quorum Sensing: Extreme Ends of a Common Spectrum. Trends Cell Biol 26:262–271.

6. Domenech A, Slager J, Veening JW. 2018. Antibiotic-Induced Cell Chaining Triggers Pneumococcal Competence by Reshaping Quorum Sensing to Autocrine-Like Signaling. Cell Rep 25:2390–2400.

7. Knoops A, Vande CF, Fontaine L, Verhaegen M, Mignolet J, Goffin P, Mahillon J, Sass A, Coenye T, Ledesma-Garcia L, Hols P. 2022. The CovRS Environmental Sensor Directly Controls the ComRS Signaling System To Orchestrate Competence Bimodality in Salivarius Streptococci. mBio 13:e0312521.

8. Chang JC, Jimenez JC, Federle MJ. 2015. Induction of a quorum sensing pathway by environmental signals enhances group A streptococcal resistance to lysozyme. Mol Microbiol 97:1097–1113.

9. Moreno-Gamez S, Sorg RA, Domenech A, Kjos M, Weissing FJ, van Doorn GS, Veening JW. 2017. Quorum sensing integrates environmental cues, cell density and cell history to control bacterial competence. Nat Commun 8:854.

10. Neiditch MB, Capodagli GC, Prehna G, Federle MJ. 2017. Genetic and Structural Analyses of RRNPP Intercellular Peptide Signaling of Gram-Positive Bacteria. Annu Rev Genet 51:311–333.

11. Fontaine L, Wahl A, Flechard M, Mignolet J, Hols P. 2015. Regulation of competence for natural transformation in streptococci. Infect Genet Evol 33:343–360.

12. Shanker E, Federle MJ. 2017. Quorum Sensing Regulation of Competence and Bacteriocins in *Streptococcus pneumoniae* and *mutans*. Genes (Basel) 8:15.

13. Fontaine L, Boutry C, de Frahan MH, Delplace B, Fremaux C, Horvath P, Boyaval P, Hols P. 2010. A novel pheromone quorum-sensing system controls the development of natural competence in *Streptococcus thermophilus* and *Streptococcus salivarius*. J Bacteriol 192:1444–1454.

14. Gardan R, Besset C, Gitton C, Guillot A, Fontaine L, Hols P, Monnet V. 2013. Extracellular life cycle of ComS, the competence-stimulating peptide of *Streptococcus thermophilus*. J Bacteriol 195:1845–1855.

15. Fontaine L, Goffin P, Dubout H, Delplace B, Baulard A, Lecat-Guillet N, Chambellon E, Gardan R, Hols P. 2013. Mechanism of competence activation by the ComRS signalling system in streptococci. Mol Microbiol 87:1113–1132.

16. Mignolet J, Fontaine L, Sass A, Nannan C, Mahillon J, Coenye T, Hols P. 2018. Circuitry Rewiring Directly Couples Competence to Predation in the Gut Dweller *Streptococcus salivarius*. Cell Rep 22:1627–1638.

17. Talagas A, Fontaine L, Ledesma-Garcia L, Mignolet J, Li de la Sierra-Gallay, Lazar N, Aumont-Nicaise M, Federle MJ, Prehna G, Hols P, Nessler S. 2016. Structural Insights into Streptococcal Competence Regulation by the Cell-to-Cell Communication System ComRS. PLoS Pathog 12:e1005980.

18. Hagen SJ, Son M. 2017. Origins of heterogeneity in *Streptococcus mutans* competence: interpreting an environment-sensitive signaling pathway. Phys Biol 14:015001.

19. Echenique J, Kadioglu A, Romao S, Andrew PW, Trombe MC. 2004. Protein serine/threonine kinase StkP positively controls virulence and competence in *Streptococcus pneumoniae*. Infect Immun 72:2434–2437.

20. Echenique JR, Chapuy-Regaud S, Trombe MC. 2000. Competence regulation by oxygen in *Streptococcus pneumoniae*: involvement of ciaRH and comCDE. Mol Microbiol 36:688–696.

21. Echenique JR, Trombe MC. 2001. Competence repression under oxygen limitation through the two-component MicAB signal-transducing system in *Streptococcus pneumoniae* and involvement of the PAS domain of MicB. J Bacteriol 183:4599–4608.

22. Pinas GE, Reinoso-Vizcaino NM, Yandar Barahona NY, Cortes PR, Duran R, Badapanda C, Rathore A, Bichara DR, Cian MB, Olivero NB, Perez DR, Echenique J. 2018. Crosstalk between the serine/threonine kinase StkP and the response regulator ComE controls the stress response and intracellular survival of *Streptococcus pneumoniae*. PLoS Pathog 14:e1007118.

23. Saskova L, Novakova L, Basler M, Branny P. 2007. Eukaryotic-type serine/threonine protein kinase StkP is a global regulator of gene expression in *Streptococcus pneumoniae*. J Bacteriol 189:4168–4179.

24. Banu LD, Conrads G, Rehrauer H, Hussain H, Allan E, van der Ploeg JR. 2010. The *Streptococcus mutans* serine/threonine kinase, PknB, regulates competence development, bacteriocin production, and cell wall metabolism. Infect Immun 78:2209–2220.

25. Dufour D, Cordova M, Cvitkovitch DG, Levesque CM. 2011. Regulation of the competence pathway as a novel role associated with a streptococcal bacteriocin. J Bacteriol 193:6552–6559.

26. Kim JN, Stanhope MJ, Burne RA. 2013. Core-gene-encoded peptide regulating virulence-associated traits in *Streptococcus mutans*. J Bacteriol 195:2912–2920.

27. Levesque CM, Mair RW, Perry JA, Lau PC, Li YH, Cvitkovitch DG. 2007. Systemic inactivation and phenotypic characterization of two-component systems in expression of *Streptococcus mutans* virulence properties. Lett Appl Microbiol 45:398–404.

28. Okinaga T, Xie Z, Niu G, Qi F, Merritt J. 2010. Examination of the *hdrRM* regulon yields insight into the competence system of *Streptococcus mutans*. Mol Oral Microbiol 25:165–177.

29. Qi F, Merritt J, Lux R, Shi W. 2004. Inactivation of the ciaH Gene in *Streptococcus mutans* diminishes mutacin production and competence development, alters sucrose-dependent biofilm formation, and reduces stress tolerance. Infect Immun 72:4895–4899.

30. Underhill SAM, Shields RC, Burne RA, Hagen SJ. 2019. Carbohydrate and PepO control bimodality in competence development by *Streptococcus mutans*. Mol Microbiol 112:1388–1402.

31. Xie Z, Okinaga T, Niu G, Qi F, Merritt J. 2010. Identification of a novel bacteriocin regulatory system in *Streptococcus mutans*. Mol Microbiol 78:1431–1447.

32. Chastanet A, Prudhomme M, Claverys JP, Msadek T. 2001. Regulation of *Streptococcus pneumoniae clp* genes and their role in competence development and stress survival. J Bacteriol 183:7295–7307.

33. Shields RC, Zeng L, Culp DJ, Burne RA. 2018. Genomewide Identification of Essential Genes and Fitness Determinants of *Streptococcus mutans* UA159. mSphere 3:e00031–18.

34. Cui L, Vigouroux A, Rousset F, Varet H, Khanna V, Bikard D. 2018. A CRISPRi screen in *E. coli* reveals sequence-specific toxicity of dCas9. Nat Commun 9:1912.

35. Liu X, Gallay C, Kjos M, Domenech A, Slager J, van Kessel SP, Knoops K, Sorg RA, Zhang JR, Veening JW. 2017. High-throughput CRISPRi phenotyping identifies new essential genes in *Streptococcus pneumoniae*. Mol Syst Biol 13:931.

36. Liu X, Kimmey JM, Matarazzo L, de B, V, Van ML, Sirard JC, Nizet V, Veening JW. 2021. Exploration of bacterial bottlenecks and *Streptococcus pneumoniae* pathogenesis by CRISPRi-Seq. Cell Host Microbe 29:107–120.

37. Van den Bogert B, Boekhorst J, Herrmann R, Smid EJ, Zoetendal EG, Kleerebezem M. 2013. Comparative genomics analysis of Streptococcus isolates from the human small intestine reveals their adaptation to a highly dynamic ecosystem. PLoS One 8:e83418.

38. Sorg RA, Kuipers OP, Veening JW. 2015. Gene expression platform for synthetic biology in the human pathogen *Streptococcus pneumoniae*. ACS Synth Biol 4:228–239.

39. Boutry C, Wahl A, Delplace B, Clippe A, Fontaine L, Hols P. 2012. Adaptor protein MecA is a negative regulator of the expression of late competence genes in *Streptococcus thermophilus*. J Bacteriol 194:1777–1788.

40. Knoops A, Ledesma-Garcia L, Waegemans A, Lamontagne M, Decat B, Degand H, Morsomme P, Soumillion P, Delvigne F, Hols P. 2022. Competence shut-off by intracellular pheromone degradation in salivarius streptococci. PLoS Genet 18:e1010198.

41. Li W, Koster J, Xu H, Chen CH, Xiao T, Liu JS, Brown M, Liu XS. 2015. Quality control, modeling, and visualization of CRISPR screens with MAGeCK-VISPR. Genome Biol 16:281.

42. Wahl A, Servais F, Drucbert AS, Foulon C, Fontaine L, Hols P. 2014. Control of natural transformation in salivarius Streptococci through specific degradation of sigmaX by the MecA-ClpCP protease complex. J Bacteriol 196:2807–2816.

43. Claverys JP, Havarstein LS. 2007. Cannibalism and fratricide: mechanisms and raisons d’etre. Nat Rev Microbiol 5:219–229.

44. Tatusov RL, Galperin MY, Natale DA, Koonin EV. 2000. The COG database: a tool for genome-scale analysis of protein functions and evolution. Nucleic Acids Res 28:33–36.

45. Neuhaus FC, Baddiley J. 2003. A continuum of anionic charge: structures and functions of D-alanyl-teichoic acids in gram-positive bacteria. Microbiol Mol Biol Rev 67:686–723.

46. Levander F, Radstrom P. 2001. Requirement for phosphoglucomutase in exopolysaccharide biosynthesis in glucose- and lactose-utilizing *Streptococcus thermophilus*. Appl Environ Microbiol 67:2734–2738.

47. Henry C, Haller L, Blein-Nicolas M, Zivy M, Canette A, Verbrugghe M, Mezange C, Boulay M, Gardan R, Samson S, Martin V, Andre-Leroux G, Monnet V. 2019. Identification of Hanks-Type Kinase PknB-Specific Targets in the *Streptococcus thermophilus* Phosphoproteome. Front Microbiol 10:1329.

48. Pereira SF, Goss L, Dworkin J. 2011. Eukaryote-like serine/threonine kinases and phosphatases in bacteria. Microbiol Mol Biol Rev 75:192–212.

49. Devine KM. 2018. Activation of the PhoPR-Mediated Response to Phosphate Limitation Is Regulated by Wall Teichoic Acid Metabolism in *Bacillus subtilis*. Front Microbiol 9:2678.

50. Eldholm V, Gutt B, Johnsborg O, Bruckner R, Maurer P, Hakenbeck R, Mascher T, Havarstein LS. 2010. The pneumococcal cell envelope stress-sensing system LiaFSR is activated by murein hydrolases and lipid II-interacting antibiotics. J Bacteriol 192:1761–1773.

51. Ganguly T, Kajfasz JK, Abranches J, Lemos JA. 2020. Regulatory circuits controlling Spx levels in *Streptococcus mutans*. Mol Microbiol 114:109–126.

52. Rojas-Tapias DF, Helmann JD. 2018. Stabilization of *Bacillus subtilis* Spx under cell wall stress requires the anti-adaptor protein YirB. PLoS Genet 14:e1007531.

53. Suntharalingam P, Senadheera MD, Mair RW, Levesque CM, Cvitkovitch DG. 2009. The LiaFSR system regulates the cell envelope stress response in *Streptococcus mutans*. J Bacteriol 191:2973–2984.

54. Ulrych A, Fabrik I, Kupcik R, Vajrychova M, Doubravova L, Branny P. 2021. Cell Wall Stress Stimulates the Activity of the Protein Kinase StkP of *Streptococcus pneumoniae*, Leading to Multiple Phosphorylation. J Mol Biol 433:167319.

55. Shah IM, Laaberki MH, Popham DL, Dworkin J. 2008. A eukaryotic-like Ser/Thr kinase signals bacteria to exit dormancy in response to peptidoglycan fragments. Cell 135:486–496.

56. Beilharz K, Novakova L, Fadda D, Branny P, Massidda O, Veening JW. 2012. Control of cell division in *Streptococcus pneumoniae* by the conserved Ser/Thr protein kinase StkP. Proc Natl Acad Sci U S A 109:E905–E913.

57. Dias R, Felix D, Canica M, Trombe MC. 2009. The highly conserved serine threonine kinase StkP of *Streptococcus pneumoniae* contributes to penicillin susceptibility independently from genes encoding penicillin-binding proteins. BMC Microbiol 9:121.

58. Fleurie A, Cluzel C, Guiral S, Freton C, Galisson F, Zanella-Cleon I, Di Guilmi AM, Grangeasse C. 2012. Mutational dissection of the S/T-kinase StkP reveals crucial roles in cell division of *Streptococcus pneumoniae*. Mol Microbiol 83:746–758.

59. Zhang R, Xu W, Shao S, Wang Q. 2021. Gene Silencing Through CRISPR Interference in Bacteria: Current Advances and Future Prospects. Front Microbiol 12:635227.

60. Tian XL, Dong G, Liu T, Gomez ZA, Wahl A, Hols P, Li YH. 2013. MecA protein acts as a negative regulator of genetic competence in *Streptococcus mutans*. J Bacteriol 195:5196–5206.

61. Shields RC, O’Brien G, Maricic N, Kesterson A, Grace M, Hagen SJ, Burne RA. 2018. Genome-Wide Screens Reveal New Gene Products That Influence Genetic Competence in *Streptococcus mutans*. J Bacteriol 200:e00508–17.

62. Dmitriev A, Mohapatra SS, Chong P, Neely M, Biswas S, Biswas I. 2011. CovR- controlled global regulation of gene expression in *Streptococcus mutans*. PLoS One 6:e20127.

63. Stevens KE, Chang D, Zwack EE, Sebert ME. 2011. Competence in *Streptococcus pneumoniae* is regulated by the rate of ribosomal decoding errors. mBio 2:e00071–11.

64. Khemici V, Prudhomme M, Polard P. 2021. Tight Interplay between Replication Stress and Competence Induction in *Streptococcus pneumoniae*. Cells 10:1938.

65. Slager J, Kjos M, Attaiech L, Veening JW. 2014. Antibiotic-induced replication stress triggers bacterial competence by increasing gene dosage near the origin. Cell 157:395–406.

66. Winther KS, Roghanian M, Gerdes K. 2018. Activation of the Stringent Response by Loading of RelA-tRNA Complexes at the Ribosomal A-Site. Mol Cell 70:95–105.

67. Kaspar J, Kim JN, Ahn SJ, Burne RA. 2016. An Essential Role for (p)ppGpp in the Integration of Stress Tolerance, Peptide Signaling, and Competence Development in *Streptococcus mutans*. Front Microbiol 7:1162.

68. Horstmann N, Saldana M, Sahasrabhojane P, Yao H, Su X, Thompson E, Koller A, Shelburne SA, III. 2014. Dual-site phosphorylation of the control of virulence regulator impacts group a streptococcal global gene expression and pathogenesis. PLoS Pathog 10:e1004088.

69. Lin WJ, Walthers D, Connelly JE, Burnside K, Jewell KA, Kenney LJ, Rajagopal L. 2009. Threonine phosphorylation prevents promoter DNA binding of the Group B Streptococcus response regulator CovR. Mol Microbiol 71:1477–1495.

70. Dufour D, Villemin C, Perry JA, Levesque CM. 2016. Escape from the competence state in *Streptococcus mutans* is governed by the bacterial population density. Mol Oral Microbiol 31:501–514.

71. Mignolet J, Fontaine L, Kleerebezem M, Hols P. 2016. Complete Genome Sequence of *Streptococcus salivarius* HSISS4, a Human Commensal Bacterium Highly Prevalent in the Digestive Tract. Genome Announc 4:e01637–15.

72. Letort C, Juillard V. 2001. Development of a minimal chemically-defined medium for the exponential growth of *Streptococcus thermophilus*. J Appl Microbiol 91:1023–1029.

73. Fontaine L, Dandoy D, Boutry C, Delplace B, de Frahan MH, Fremaux C, Horvath P, Boyaval P, Hols P. 2010. Development of a versatile procedure based on natural transformation for marker-free targeted genetic modification in *Streptococcus thermophilus*. Appl Environ Microbiol 76:7870–7877.

74. Dorrazehi GM, Worms S, Chirakadavil JB, Mignolet J, Hols P, Soumillion S. 2020. Building Scarless Gene Libraries in the Chromosome of Bacteria, p 189-211. *In*: Iranzo O, Roque A (ed), Peptide and Protein Engineering, Springer Protocols Handbooks. Humana, New York, NY.

75. Lu S, Wang J, Chitsaz F, Derbyshire MK, Geer RC, Gonzales NR, Gwadz M, Hurwitz DI, Marchler GH, Song JS, Thanki N, Yamashita RA, Yang M, Zhang D, Zheng C, Lanczycki CJ, Marchler-Bauer A. 2020. CDD/SPARCLE: the conserved domain database in 2020. Nucleic Acids Res 48:D265–D268.

76. Ogura M, Hashimoto H, Tanaka T. 2002. Med, a cell-surface localized protein regulating a competence transcription factor gene, *comK*, in *Bacillus subtilis*. Biosci Biotechnol Biochem 66:892–896.

