## Supplemental Fig S1 for "A genome-wide CRISPRi screen reveals a StkP-mediated connection between cell-wall integrity and competence in *Streptococcus salivarius*"

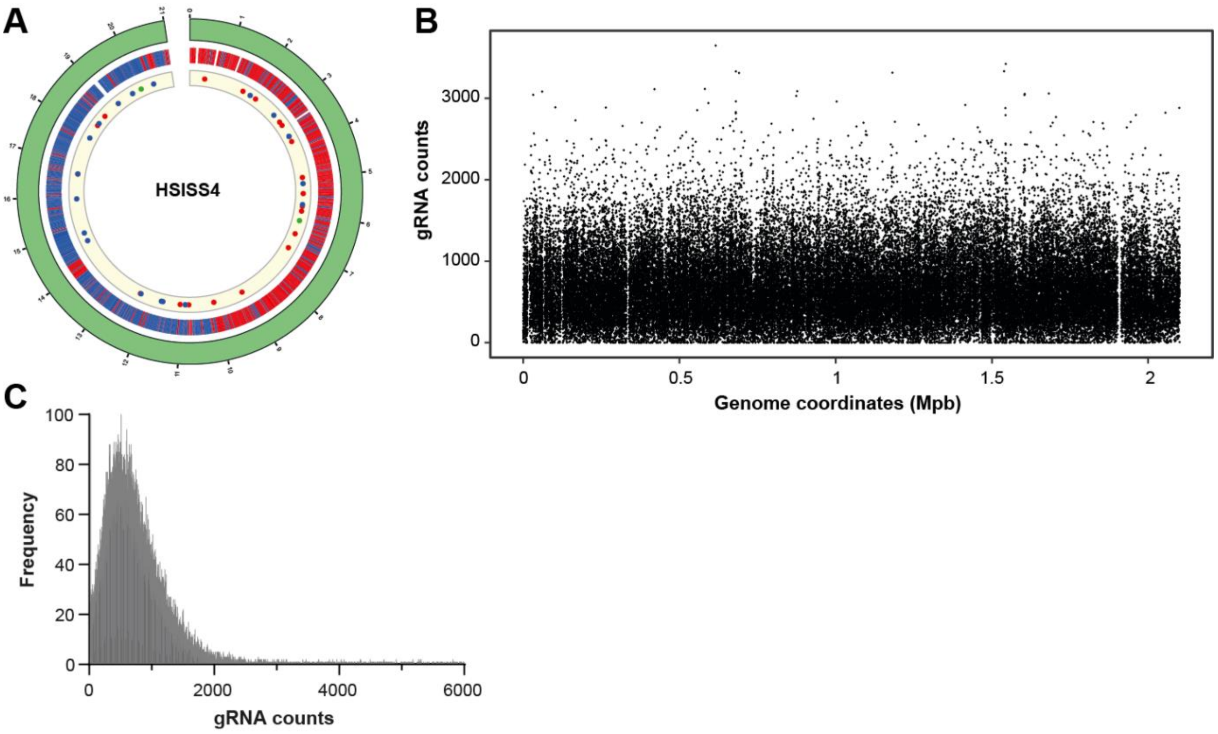


**Figure S1. Random chromosomal distribution of the gRNA library. A.** 40 randomly picked colonies from the gRNA library were PCR-amplified, Sanger-sequenced and mapped on the *Streptococcus salivarius* HSISS4 genome. From outside to center: numbers denote the genomic position (× $10^5$ bp), red and blue regions depict coding strand being on (+) or (-) strand, respectively. Large color-empty regions correspond to clusters of tRNAs or rRNAs. Red and blue dots show the mapping of gRNAs targeting the (+) or (-) strands, respectively. Green dots show gRNAs targeting intergenic regions. **B.** NGS mapping of the reads from gRNAs in the mock library (no library induction). The numbers of reads per gRNA are shown. gRNAs with common sequences were discarded since their mapping at multiple sites biases the analysis. Low density mapping of gRNAs on the graph are associated to highly similar sequences such as rRNA, tRNA or multiple insertion of transposons. Removal of gRNAs with the same sequences from the analysis particularly influences the mapping in those regions. **C.** Frequency distribution of gRNA counts from the mock library.
