## Supplemental Fig S2 for "A genome-wide CRISPRi screen reveals a StkP-mediated connection between cell-wall integrity and competence in *Streptococcus salivarius*"

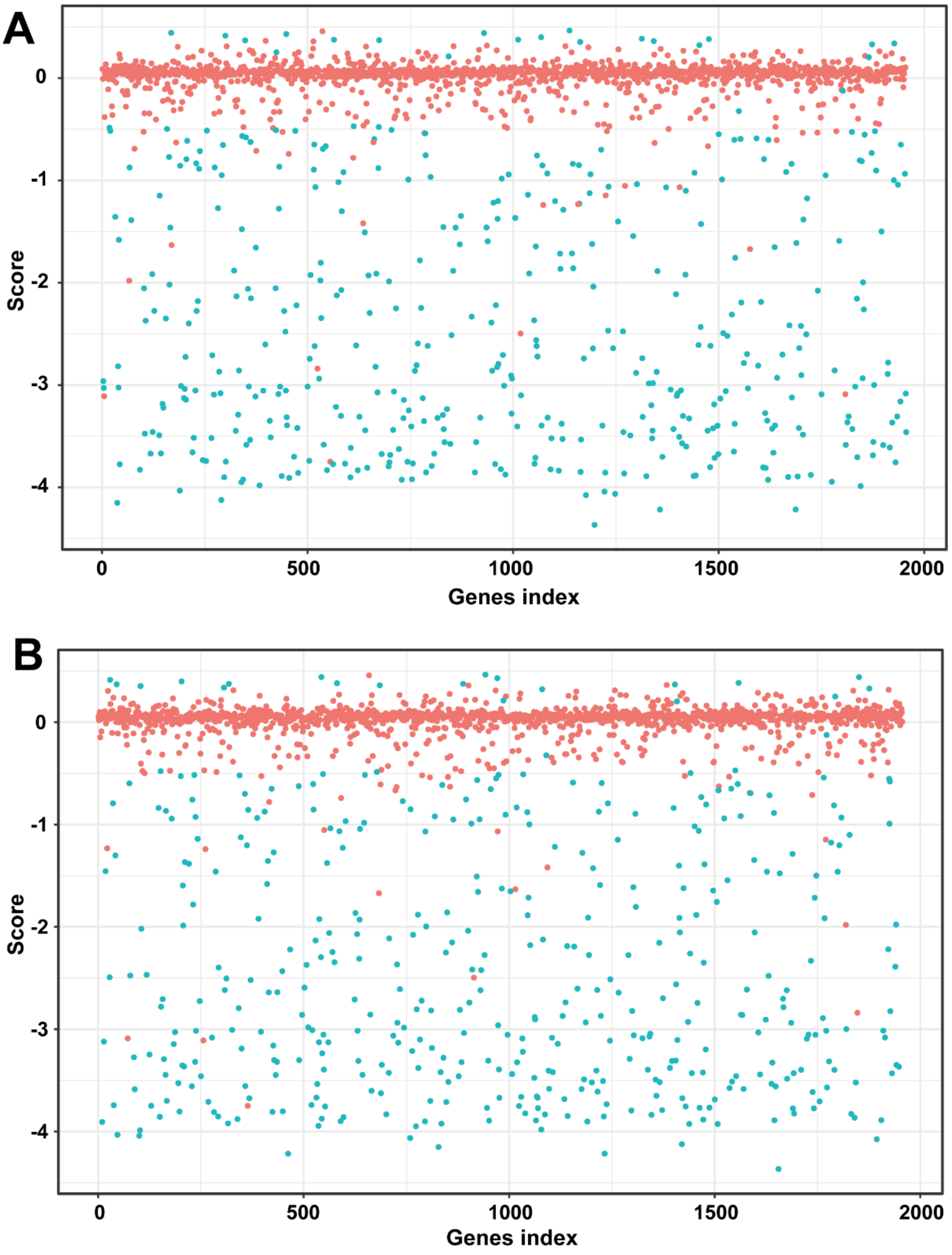


**Figure S2. Gene-associated gRNA depletion scores.** gRNA reads computed with library induction (**A**) and with library induction plus competence activation (**B**) were both compared to the reads computed in the mock condition thanks to the MAGeCK algorithm (1). The algorithm generated a score translating the total depletion of gRNAs for one gene and a False Discovery Rate value (FDR), as a significative marker of the score. The plots show the score computed for each gene (red, FDR>0.05; blue, FDR<0.05). Each gene was associated to a random number (gene index) for the sake of clarity. Values for each gene can be found in Tables S2 and S3.

### REFERENCE

1. Li W, Koster J, Xu H, Chen CH, Xiao T, Liu JS, Brown M, Liu XS. 2015. Quality control, modeling, and visualization of CRISPR screens with MAGeCK-VISPR. *Genome Biol* 16:281.
