## Supplemental Fig S3 for "A genome-wide CRISPRi screen reveals a StkP-mediated connection between cell-wall integrity and competence in *Streptococcus salivarius*"

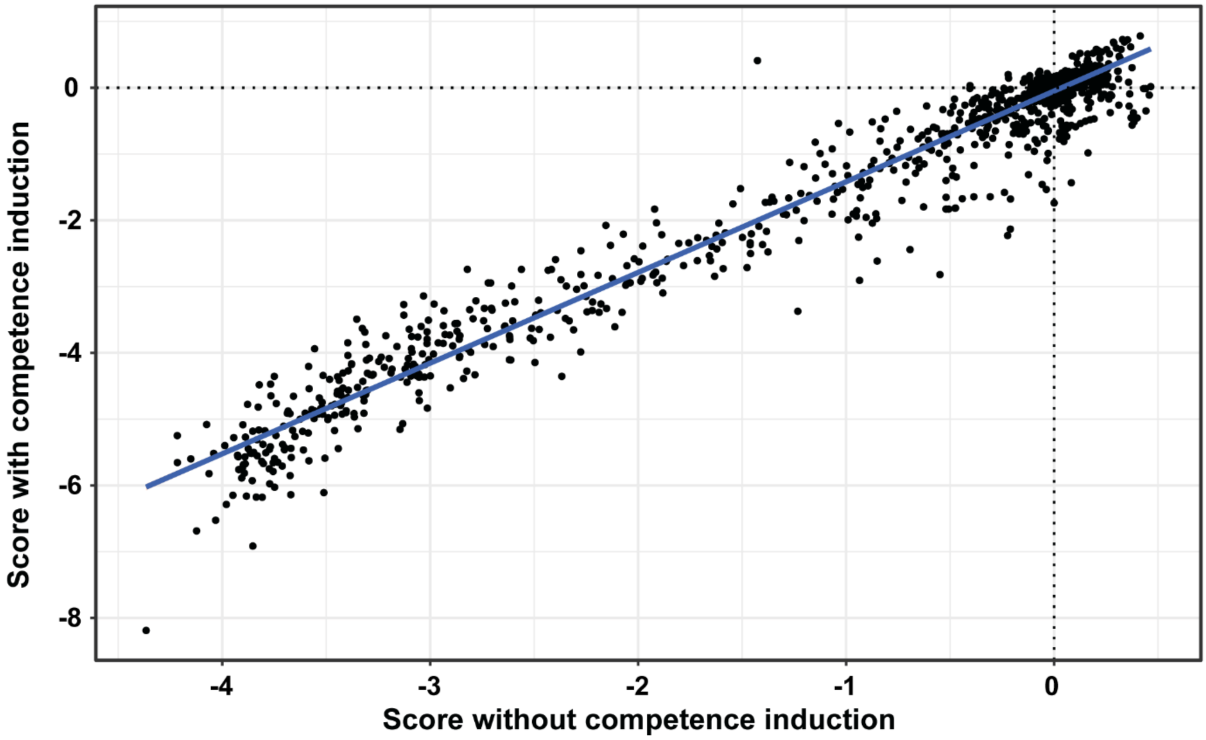


**Figure S3. Linear regression of gene-associated gRNA depletion scores.** gRNA depletion for each gene in comparison to the mock condition (Fig. S2) are plotted against each other. The linear regression was computed thanks to the *lm* function from R using the QR method ($R^2$=0.97). Dashed lines denote a score of zero associated to a neutral fitness effect of gene inhibition.
