## Supplemental Table S7 for "A genome-wide CRISPRi screen reveals a StkP-mediated connection between cell-wall integrity and competence in *Streptococcus salivarius*"

**Table S7A. List of bacterial strains used in this study**

| Strain | Characteristics | Reference/source |
|---|---|---|
| ***Streptococcus salivarius*** | | |
| HSISS4 | Wild-type gastro-intestinal tract isolate | (1) |
| AK0004 | HSISS4 *tRNA*$_{thr}$::P$_{comX}$-*luxAB-spec*<br>*tRNA*$_{ser}$::P$_{xyl2}$-*comR-cat* | (2) |
| AK0033 | HSISS4 *tRNA*$_{thr}$::P$_{F6}$-*lacI-lox72*<br>*tRNA*$_{ser}$::P$_{lac}$-*dcas9-lox72*<br>*tnpII*::*P*$_{comX}$-*luxAB-lox72* | (2) |
| AK0034 | HSISS4 *tRNA*$_{thr}$::P$_{F6}$-*lacI-lox72*<br>*tRNA*$_{ser}$::P$_{lac}$-*dcas9-lox72*<br>*tnpII*::*P*$_{comR}$-*luxAB-lox72* | (2) |
| AK0046 | HSISS4 *tRNA*$_{thr}$::P$_{F6}$-*lacI-lox72*<br>*tRNA*$_{ser}$::P$_{lac}$-*dcas9-lox72*<br>*tnpII*::*P*$_{comX}$-*luxAB-lox72*<br>*SUC*::*P*$_{xyl2}$-*comR-lox72* | (3) |
| AK0065 | HSISS4 *tRNA*$_{thr}$::P$_{F6}$-*lacI-lox72*<br>*tRNA*$_{ser}$::P$_{lac}$-*dcas9-lox72*<br>*tnpII*::*P*$_{comX}$-*luxAB-lox72*<br>*SUC*::*P*$_{xyl2}$-*comR-cat*<br>*P*$_{comX}$-*lacZ-spc* | This work |
| AK0066 | HSISS4 *tRNA*$_{thr}$::P$_{F6}$-*lacI-lox72*<br>*tRNA*$_{ser}$::P$_{lac}$-*dcas9-lox72*<br>*tnpII*::*P*$_{comX}$-*luxAB-lox72*<br>*SUC*::*P*$_{xyl2}$-*comR-lox72*<br>*GOR*::*P*$_{3}$-*g_30(HSISS4_01622)-erm* | This work |
| AK0067 | HSISS4 *tRNA*$_{thr}$::P$_{F6}$-*lacI-lox72*<br>*tRNA*$_{ser}$::P$_{lac}$-*dcas9-lox72*<br>*tnpII*::*P*$_{comX}$-*luxAB-lox72*<br>*SUC*::*P*$_{xyl2}$-*comR-lox72*<br>*GOR*::*P*$_{3}$-*g_27(HSISS4_01391)-erm* | This work |
| AK0068 | HSISS4 *tRNA*$_{thr}$::P$_{F6}$-*lacI-lox72*<br>*tRNA*$_{ser}$::P$_{lac}$-*dcas9-lox72*<br>*tnpII*::*P*$_{comX}$-*luxAB-lox72*<br>*SUC*::*P*$_{xyl2}$-*comR-lox72*<br>*GOR*::*P*$_{3}$-*g_31(HSISS4_00663)-erm* | This work |
| AK0069 | HSISS4 *tRNA*$_{thr}$::P$_{F6}$-*lacI-lox72*<br>*tRNA*$_{ser}$::P$_{lac}$-*dcas9-lox72*<br>*tnpII*::*P*$_{comX}$-*luxAB-lox72*<br>*SUC*::*P*$_{xyl2}$-*comR-lox72*<br>*GOR*::*P*$_{3}$-*g_32(HSISS4_00805)-erm* | This work |
| AK0070 | HSISS4 *tRNA*$_{thr}$::P$_{F6}$-*lacI-lox72*<br>*tRNA*$_{ser}$::P$_{lac}$-*dcas9-lox72*<br>*tnpII*::*P*$_{comX}$-*luxAB-lox72* | This work |

| | | |
|---|---|---|
| | *SUC*::*P*$_{xyl2}$-*comR-lox72*<br>*GOR*::*P*$_{3}$-*g_35(HSISS4_01302)-erm* | |
| AK0071 | HSISS4 *tRNA*$_{thr}$::P$_{F6}$-*lacI-lox72*<br>*tRNA*$_{ser}$::P$_{lac}$-*dcas9-lox72*<br>*tnpII*::*P*$_{comX}$-*luxAB-lox72*<br>*SUC*::*P*$_{xyl2}$-*comR-lox72*<br>*GOR*::*P*$_{3}$-*g_26(gpmB-dacB-mur3)-erm* | This work |
| AK0072 | HSISS4 *tRNA*$_{thr}$::P$_{F6}$-*lacI-lox72*<br>*tRNA*$_{ser}$::P$_{lac}$-*dcas9-lox72*<br>*tnpII*::*P*$_{comX}$-*luxAB-lox72*<br>*SUC*::*P*$_{xyl2}$-*comR-lox72*<br>*GOR*::*P*$_{3}$-*g_37(clpC)-erm* | This work |
| AK0073 | HSISS4 *tRNA*$_{thr}$::P$_{F6}$-*lacI-lox72*<br>*tRNA*$_{ser}$::P$_{lac}$-*dcas9-lox72*<br>*tnpII*::*P*$_{comX}$-*luxAB-lox72*<br>*SUC*::*P*$_{xyl2}$-*comR-lox72*<br>*GOR*::*P*$_{3}$-*g_38(clpC)-erm* | This work |
| AK0074 | HSISS4 *tRNA*$_{thr}$::P$_{F6}$-*lacI-lox72*<br>*tRNA*$_{ser}$::P$_{lac}$-*dcas9-lox72*<br>*tnpII*::*P*$_{comX}$-*luxAB-lox72*<br>*SUC*::*P*$_{xyl2}$-*comR-lox72*<br>*GOR*::*P*$_{3}$-*g_39(pepF)-erm* | This work |
| AK0075 | HSISS4 *tRNA*$_{thr}$::P$_{F6}$-*lacI-lox72*<br>*tRNA*$_{ser}$::P$_{lac}$-*dcas9-lox72*<br>*tnpII*::*P*$_{comX}$-*luxAB-lox72*<br>*SUC*::*P*$_{xyl2}$-*comR-lox72*<br>*GOR*::*P*$_{3}$-*g_40(scuR/sarF)-erm* | This work |
| AK0076 | HSISS4 *tRNA*$_{thr}$::P$_{F6}$-*lacI-lox72*<br>*tRNA*$_{ser}$::P$_{lac}$-*dcas9-lox72*<br>*tnpII*::*P*$_{comX}$-*luxAB-lox72*<br>*SUC*::*P*$_{xyl2}$-*comR-lox72*<br>*GOR::P3-g_41(pepXP)-erm* | This work |
| AK0077 | HSISS4 *tRNA*$_{thr}$::P$_{F6}$-*lacI-lox72*<br>*tRNA*$_{ser}$::P$_{lac}$-*dcas9-lox72*<br>*tnpII*::*P*$_{comX}$-*luxAB-lox72*<br>*SUC*::*P*$_{xyl2}$-*comR-lox72*<br>*GOR*::*P*$_{3}$-*g_42(carB)-erm* | This work |
| AK0078 | HSISS4 *tRNA*$_{thr}$::P$_{F6}$-*lacI-lox72*<br>*tRNA*$_{ser}$::P$_{lac}$-*dcas9-lox72*<br>*tnpII*::*P*$_{comX}$-*luxAB-lox72*<br>*SUC*::*P*$_{xyl2}$-*comR-lox72*<br>*GOR*::*P*$_{3}$-*g_43(IG664598)-erm* | This work |
| AK0079 | HSISS4 *tRNA*$_{thr}$::P$_{F6}$-*lacI-lox72*<br>*tRNA*$_{ser}$::P$_{lac}$-*dcas9-lox72* | This work |

| | | |
|---|---|---|
| | *tnpII::*$P_{comR}$*-luxAB-lox72* | |
| | *GOR::*$P_3$*-g_23(stkP)-erm* | |
| AK0080 | HSISS4 *tRNA*$_{thr}$*::*$P_{F6}$*-lacI-lox72*<br>*tRNA*$_{ser}$*::*$P_{lac}$*-dcas9-lox72*<br>*tnpII::*$P_{comX}$*-luxAB-lox72*<br>*SUC::*$P_{xyl2}$*-comR-lox72*<br>*GOR::*$P_3$*-g_23(stkP)-erm* | This work |
| ***Bacillus subtilis*** | | |
| 168 | Wild-type | J. Mahillon, Laboratory collection |

**Table S7B. List of plasmids used in this study**

| Plasmid | Characteristic(s) | Reference |
|---|---|---|
| pGhost-cre | Thermosensitive replication origin vector in *S. salivarius*, encoding the Cre recombinase; $ery^R$ | (4) |
| pGIUD0855ery | pUC18 derivative containing the *erm* gene | (4) |
| pJUD*specmut1-gfp*$^{+}$ter | Terminator associated-*gfp*$^{+}$ ORF cloned in cloned in pJUDspecmut1 | (5) |
| pJIM*cat* | pJIM4900 derivative with a *cat* cassette | (5) |

**Table S7C. List of oligonucleotides used in this study**

| Name | Sequence (5' to 3') |
|---|---|
| **AK396** | TCAACCTCCTATTAATAGATATAATTTTTG |
| **AK458** | GTAGCAACACTCTTGTTTAAAG |
| **AK459** | GTGGCTGAATTATCAAAATAAATC |
| **AK465** | TATAGATTTCATTGCTGGAC |
| **AK484** | GCCTTAGCCAAATCGTAATC |
| **AK485** | AACAGGAGGTTTTCGTAATGG |
| **AK518** | GTATTCAAAAAACAACTAACAGG |
| **AK519** | CCAACGTCCAGCAATGAAATCTATATAAGGAAGATAAATCCCATAAGG |
| **AK520** | GTTATTACCTTCAAAAAAGGAGAATAATCTTTCACGTTACTAAAGGGAATGTA |
| **Dn.Rv.lox71** | TTCACGTTACTAAAGGGAATGTA |
| **ML7** | GGAGGCAAAGTCCAATAATACTTAAGGAAGATAAATCCCATAAG |
| **ML8** | GTTATTACCTTCAAAAAAGGAGAATAATCTTTCACGTTACTAAAGGGAATG |
| **ML9** | AGATTATTCTCCTTTTTTGAAG |
| **ML11** | GTTTGAATTTTTTCAGTCGTGTTCATTCAACCTCCTATTAATAGATATAATTTTTG |
| **ML13** | TTAGTTGAGTGGTTCAATCATG |
| **Up.Fw.lox66** | TAAGGAAGATAAATCCCATAAGG |

**Table S7D. List of PCR fragments used in this study**

| Strain/DNA | PCR fragment | Template DNA | Primer 1 | Primer 2 |
|---|---|---|---|---|
| AK0065 | Up HR of *lacZ* locus | HSISS4 | ML13 | ML11 |
| | Promoter of *comX*, $P_{comX}$ | HSISS4 | AK396 | AK465 |
| | Spectinomycin resistance cassette, *spc* | pJUD*specmut1-gfp*+ter | AK519 | AK520 |
| | Dw HR of *lacZ* locus | HSISS4 | ML9 | AK518 |
| AK0066 | $P_3$-g_30(*HSISS4_01622*) fused to the two HR of the *GOR* locus together with *erm* | Selected colony from transformation screen #2 | AK458 | AK459 |
| AK0067 | $P_3$-g_27(*HSISS4_01391*) fused to the two HR of the GOR locus together with *erm* | Selected colony from transformation screen #5 | AK458 | AK459 |
| AK0068 | $P_3$-g_31(*HSISS4_00663*) fused to the two HR of the GOR locus together with *erm* | Selected colony from transformation screen #6 | AK458 | AK459 |
| AK0069 | $P_3$-g_32(*HSISS4_00805*) fused to the two HR of the GOR locus together with *erm* | Selected colony from transformation screen #9 | AK458 | AK459 |
| AK0070 | $P_3$-g_35(*HSISS4_01302*) fused to the two HR of the GOR locus together with *erm* | Selected colony from transformation screen #13 | AK458 | AK459 |
| AK0071 | $P_3$-g_26(*gpmB-dacB-mur3*) fused to the two HR of the GOR locus together with *erm* | Selected colony from transformation screen #16 | AK458 | AK459 |
| AK0072 | $P_3$-g_37(*clpC*) fused to the two HR of the GOR locus together with *erm* | Selected colony from transformation screen #17 | AK458 | AK459 |
| AK0073 | $P_3$-g_38(*clpC*) fused to the two HR of the GOR locus together with *erm* | Selected colony from transformation screen #18 | AK458 | AK459 |
| AK0074 | $P_3$-g_39(*pepF*) fused to the two HR of the GOR locus together with *erm* | Selected colony from transformation screen #21 | AK458 | AK459 |
| AK0075 | $P_3$-g_40(*scuR/sarF*) fused to the two HR of the GOR locus together with *erm* | Selected colony from transformation screen #22 | AK458 | AK459 |
| AK0076 | $P_3$-g_41(*pepXP*) fused to the two HR of the GOR locus together with *erm* | Selected colony from transformation screen #23 | AK458 | AK459 |
| AK0077 | $P_3$-g_42(*carB*) fused to the two HR of the GOR locus together with *erm* | Selected colony from transformation screen #25 | AK458 | AK459 |
| AK0078 | $P_3$-g_43(*IG664598*) fused to the two HR of the GOR locus together with *erm* | Selected colony from transformation screen #28 | AK458 | AK459 |
| AK0079, AK0080 | $P_3$-g_27(*stkP*) fused to the two HR of the GOR locus together with *erm* | Selected colony from the *lacZ* screen | AK458 | AK459 |
| Δ*lacZ::cat* | Up HR of *lacZ* locus (2000 bp) | HSISS4 | AK484 | ML7 |
| | Chloramphenicol resistance cassette, *cat* | pJIM*cat* | Up.Fw.lox66 | Dn.Rv.lox71 |
| | Dw HR of *lacZ* locus (2000 bp) | HSISS4 | ML8 | AK485 |

### REFERENCES


1. Van den Bogert B, Boekhorst J, Herrmann R, Smid EJ, Zoetendal EG, Kleerebezem M. 2013. Comparative genomics analysis of Streptococcus isolates from the human small intestine reveals their adaptation to a highly dynamic ecosystem. *PLoS One* 8:e83418.

2. Knoops A, Vande CF, Fontaine L, Verhaegen M, Mignolet J, Goffin P, Mahillon J, Sass A, Coenye T, Ledesma-Garcia L, Hols P. 2022. The CovRS Environmental Sensor Directly Controls the ComRS Signaling System To Orchestrate Competence Bimodality in Salivarius Streptococci. *mBio* 13:e0312521.

3. Knoops A, Ledesma-Garcia L, Waegemans A, Lamontagne M, Decat B, Degand H, Morsomme P, Soumillion P, Delvigne F, Hols P. 2022. Competence shut-off by intracellular pheromone degradation in salivarius streptococci. *PLoS Genet* 18:e1010198.

4. Fontaine L, Dandoy D, Boutry C, Delplace B, de Frahan MH, Fremaux C, Horvath P, Boyaval P, Hols P. 2010. Development of a versatile procedure based on natural transformation for marker-free targeted genetic modification in *Streptococcus thermophilus*. *Appl Environ Microbiol* 76:7870-7877.

5. Mignolet J, Fontaine L, Sass A, Nannan C, Mahillon J, Coenye T, Hols P. 2018. Circuitry Rewiring Directly Couples Competence to Predation in the Gut Dweller *Streptococcus salivarius*. *Cell Rep* 22:1627-1638.
